# Evolutionary persistence of a highly prevalent multicopy mitochondrial-derived nuclear insertion (Mega-NUMT) in Neotropical *Drosophila* flies

**DOI:** 10.64898/2026.03.31.715258

**Authors:** Merce Montoliu-Nerin, Anton Strunov, Eleanor Heyworth, Daniela I. Schneider, Julia Thoma, Aurélie Hua-Van, Cécile Courret, Lisa Klasson, Wolfgang J. Miller

## Abstract

Although strict maternal transmission of mitochondria is a general feature of animals for ensuring homogeneity in mitochondrial DNA (mtDNA) across generations, exceptions were reported in the recent past. For example, some extremely rare but spectacular cases of heteroplasmy and paternal transmission in humans have questioned the universal evolutionary principle. Hence, as an alternative, the Mega-NUMT concept was coined to explain this discovery and was thereafter partly proven to exist. This concept expands on the quite common transfer of mtDNA fragments to the nucleus (NUMTs) by considering the existence of multicopy mitochondrial nuclear insertions. Mega-NUMT reports are currently restricted to a few cases in animals, including humans. However, their detailed genomic organization, natural prevalence, and potential biological functions remain unclear.

Here, we discovered that up to 60 full-sized mitochondrial genomes are integrated into the nuclear genome of the neotropical *Drosophila paulistorum* using long-read sequencing and *in situ* hybridization. The copies are organized in one cluster on chromosome 3, which we designated the “Dpau Mega-NUMT”. Contrary to the rarity in humans, this Mega-NUMT is found at high prevalence (40%) in both laboratory lines and natural *D. paulistorum* populations of different semispecies. Additionally, the Mega-NUMT copies are phylogenetically separated from the current mitotypes of *D. paulistorum*. Together, these observations suggest long-term maintenance of the Mega-NUMT in nature. Hence, we propose that the Dpau Mega-NUMT may have been transferred to the nuclear genome before the *D. paulistorum* semispecies radiation and speculate based on these findings that it possibly is maintained at relatively high prevalence in nature by balancing selection or due to a yet undetermined function.

## Introduction

Mitochondrial DNA (mtDNA) has been widely used as a genetic marker for molecular systematics of animals for more than 30 years (Harrison, 1989). The high copy number of mtDNA per cell, as well as its rapid mutation rate, makes it particularly informative for investigating phylogenetic relationships and evolutionary history (Brown et al., 1979). Mitochondrial markers have been extensively utilized to resolve taxonomic uncertainties and to identify cryptic species (Hebert et al., 2003; Elías-Gutiérrez et al., 2008). Additionally, the use of mitochondrial data allows the reconstruction of phylogeographic patterns, providing insights into historical population dynamics and speciation events (Avise et al., 1983; Baião et al., 2023).

Studies from the pre-genomic era using Southern blots and PCR have revealed the existence of nuclear mitochondrial DNA segments (NUMTs), described as fragments of mtDNA that have become integrated into the nuclear genome (Lopez et al., 1994; Hazkani-Covo & Graur, 2007). Although the mechanisms of integration are not yet fully described, a common explanation outlines the insertion of mitochondrial sequences during the repair of double-strand chromosomal breaks (Blanchard & Schmidt, 1995, 1996; Bensasson et al., 2001). NUMTs have primarily been found in genomic regions with high levels of repetitive elements (Behura, 2007; Tsuji et al., 2012; Dayama et al., 2014; Schiavo et al., 2017; Wang et al., 2020) and distinctive GC content (Tsuji et al., 2012; Dayama et al., 2014; Schiavo et al., 2017). After their integration, NUMTs often degrade quickly, as the different genetic codes in the nuclear and mitochondrial genomes prevent proper translation of the genes (Bensasson et al., 2001). Nonetheless, some functions associated with NUMTs have been found, including the formation of new exons (Noutsos et al., 2007) and regulatory elements (Vendrami et al., 2022).

The presence of NUMTs can lead to misinterpretations when studying mitochondrial genomes (Calvignac et al., 2011), as it can result in erroneous inferences of relationships between lineages (Kress et al., 2015; DeSalle & Goldstein, 2019). Furthermore, the presence of NUMTs may lead to wrongful inference of heteroplasmy (Parr et al., 2006; Balciuniene & Balciunas, 2019), which is the coexistence of more than one mtDNA sequence variant within an individual. The misinterpretations arise from the difficulty in distinguishing NUMTs from real heteroplasmic variants when using techniques such as PCR. However, the high copy number of mtDNA compared to nuclear loci containing NUMTs may commonly suffice to mask the NUMT signal. Even so, one must be cautious (Parfait et al., 1998), as several studies have reviewed investigations in which high copy numbers of NUMTs in the genome had been interpreted as heteroplasmy (Hirano et al., 1997; Salas et al., 2005; Yao et al., 2006; Lutz-Bonengel et al., 2021). To describe such multicopy NUMTs, the term Mega-NUMT was coined as a theoretical concept in response to a claim of paternal transmission of mitochondria in seemingly heteroplasmic humans (Balciuniene & Balciunas, 2019; Lutz-Bonengel et al., 2021). However, large multicopy NUMTs, covering approximately half of the mt-genome and with an estimated 38-76 copies, had also previously been found in the domestic cat and its close relatives (Lopez et al., 1994) and later also identified in several cat sequences using long-read sequencing, albeit in somewhat fewer copies (Patterson et al., 2023).

In this study, we focus on the Neotropical fly *Drosophila paulistorum* spp., commonly used as a study system for the evolutionary process of sympatric speciation (Burla et al., 1949; Dobzhansky & Spassky, 1959; Dobzhansky et al., 1964). The *D. paulistorum* superspecies belongs to the D. willistoni group and is composed of six semispecies, named Amazonian (AM), Andean Brazilian (AB), Centro-American (CA), Interior (IN), Orinocan (OR), and Transitional (TR), presenting different levels of pre- and post-mating incompatibilities (Dobzhansky & Spassky, 1959). Phylogenetic incongruences between the nuclear and mitochondrial DNA of *D. paulistorum* spp. have been described (Gleason et al., 1998; Robe et al., 2010; Baião et al., 2023), suggesting past introgression events. Additionally, a recent phylogenomic analysis of *D. paulistorum* spp. revealed that their mitochondrial genomes are polyphyletic and split into two major clades, α and β (Baião et al., 2023).

Here, we identify a *D. paulistorum* line from the OR semispecies, O11, with an apparently fixed presence of the two different mitotypes, α and β. However, our analysis revealed that in addition to the presence of a β mitochondrion, O11 has a NUMT consisting of around 60 copies of nearly complete mt-genomes with high similarity to the α mitotype on chromosome 3. We designate this nuclear insertion as the *Dpau* Mega-NUMT. Furthermore, we found that this Mega-NUMT is present in at least one line from three of the six described *D. paulistorum* semispecies and has a relatively high prevalence of around 40% in fly populations from French Guiana. Taken together, these data suggest that the *Dpau* Mega-NUMT was present before or just after the split of the semispecies and may be maintained in nature by mechanisms like balancing selection or others that are yet still undetermined.

## Results

### The two mitotypes, α and β, are fixed in individual flies of the *D. paulistorum* line O11

We sequenced PCR products of the mitochondrial *Cytochrome Oxidase I* (*COI*) gene from several *D. paulistorum* lines for mt-barcoding. We found multiple robust double peaks in the resulting chromatograms derived from single flies of the OR line O11, suggesting the presence of both α and β mitotypes within the same individual. The presence of a *Hin*dIII restriction site for the α-mitotype but not ß (**Fig. 1A**) was used to perform diagnostic RFLP/*Hin*dIII Southern blots on DNA from single O11 flies (Junakovic, 2004) probed with the *COI* cloned plasmid (**Fig. 1B** and **Table S1, Fig. S1A**). We observed that the O11 line again showed a clear heteroplasmic pattern, with both α and β mitotypes represented in individual flies at approximately similar abundances, suggesting almost equal copy numbers (**Fig. 1B**). In contrast, single flies of line C2 belonging to the CA semispecies and line A28 from the AM semispecies were homoplasmic and carried either the α or the β mitotype, respectively (**Fig. 1B**). Furthermore, we performed *CO1*-specific RFLP/*Hin*dIII-PCR on multiple single females and males (**Table S1, Fig. S1**) which showed a consistency on the presence of both α and β mitotypes in all tested O11 flies (**Fig. 1C**), which suggested fixation in this long-term lab line collected in the 1960s.

**Figure 1.**
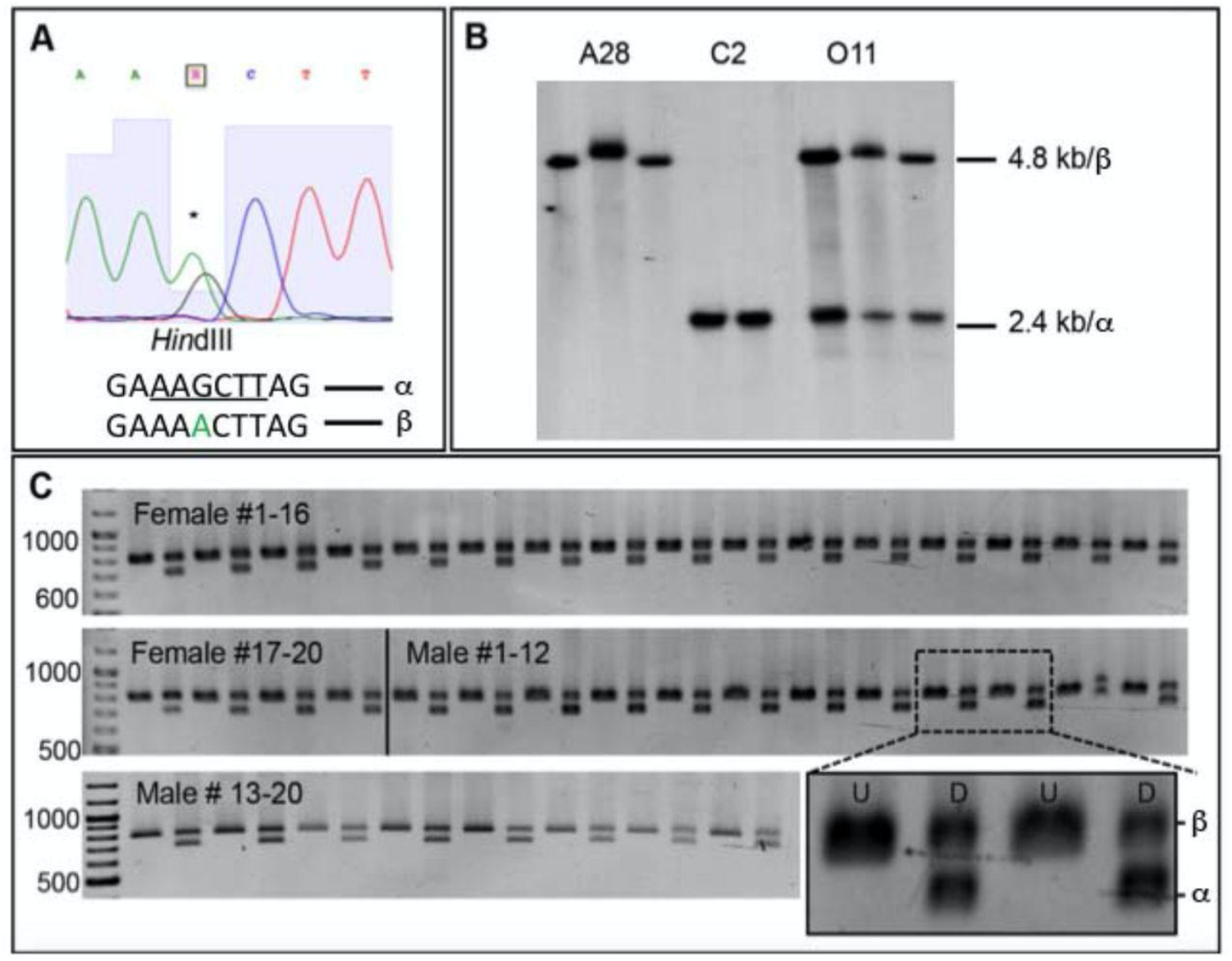
Presence of two mitotypes in the *D. paulistorum* Orinocan line O11. (**A**) Chromatogram derived from *COI* direct Sanger sequencing of one O11 female covering the α-diagnostic *Hin*dIII restriction site (underlined). The asterisk highlights the double peak with Guanine (G) for the α- and Adenine (A) for the β-mitotype. (**B**) Single-fly *CO1* Southern blot digested with *Hin*dIII and probed with cloned *CO1*. While the Amazonian females of the A28 line (n=3) present only the β-mitotype (4.8kb) and the females of the Centro-American line C2 (n=2) only the α-mitotype (2.4kb), all randomly picked Orinocan O11 females (n=3) present both the α- and β-mitotypes (4.8 and 2.4kb). (**C**) α-diagnostic *CO1* RFLP/*Hin*dIII-PCR screen (**Fig. S1 A-C**) on O11 single flies (n=20 for each sex). Undigested (U) and digested (D) CO1-PCR fragments per fly are shown exemplary for two individual specimens (magnification at the bottom right in C). All tested single flies (40/40) exhibited clear double bands after *Hin*dIII digestion.

Subsequently, we measured the copy numbers of both α and β mitochondrial (mt) genomes, relative to the nuclear genome, using discriminant qPCR on the *CO3* gene (see Materials and Methods). The quantification of both the α and β mt genomes was in multiple tissues of both females and males and resulted in similar average values of 27.2±13.9 α and 31.6±22.7 β mt genome copies per nuclear genome (**Fig. 2**). The β mt-genome copy numbers were strongly affected by tissue (**Table S2**; df=4, F-value=27.694, p<0.001, F-test, **Fig. 2A**, **Table S2**) and a more subtly, yet significantly, by age and sex combined (**Table S2**; df=1, F-value=6.66, p<0.05, F-test), and by sex alone (**Table S2**; df=1, F-value=4.714, p<0.05, F-test), as expected based on mitochondrial function. In contrast, the number of copies of the α mt-genome was not significantly affected by any of the tested parameters (**Fig. 2B**, **Table S2**), suggesting that the α mitotype is not a typical mitochondrion.

**Figure 2.**
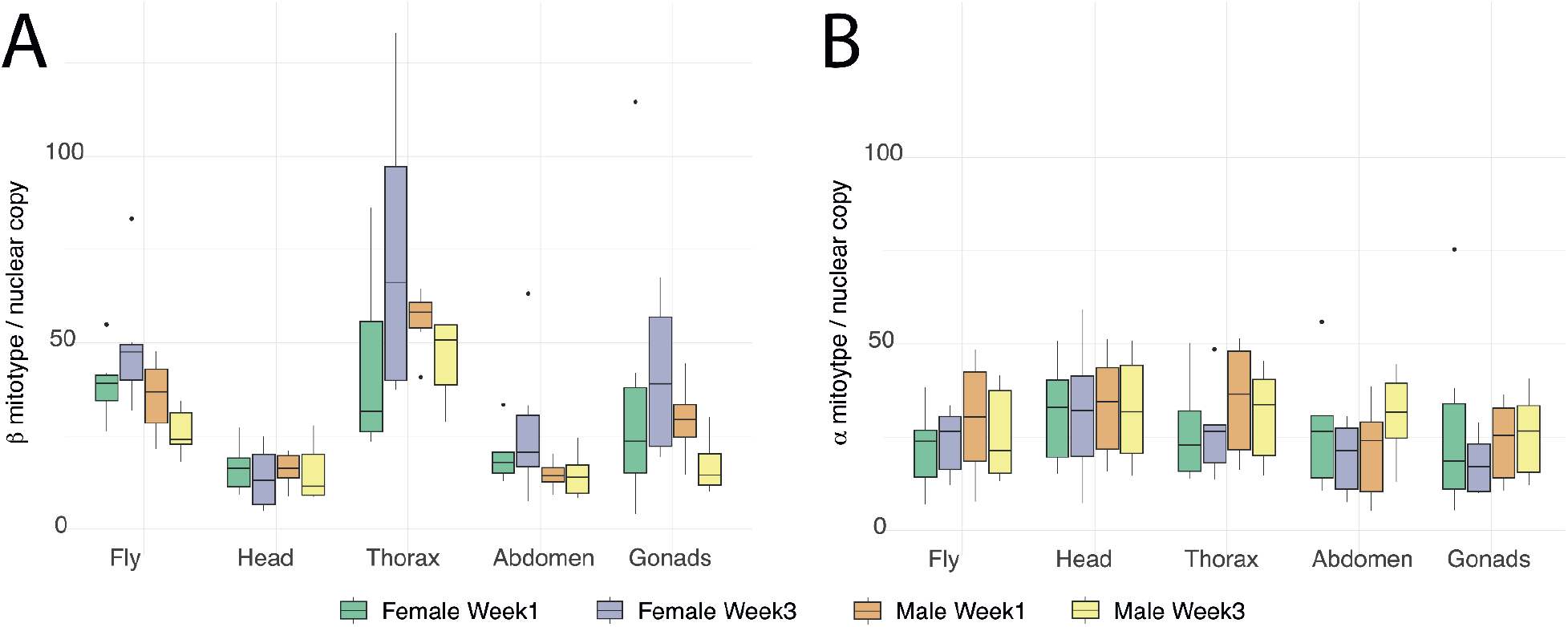
qPCR quantification of α and β mitotypes in *D. paulistorum* O11 flies. 120 flies were analyzed to assess the number of α and β mitotype copies relative to the nuclear genome copies using discriminant qPCR. Boxplots show the distribution of values, mean, and standard deviation for (**A**) the β mitotype and (**B**) the α mitotype.

### Discovery of a Mega-NUMT with long reads

Given the seemingly fixed high abundance of two mitotypes in one individual and unresponsive behavior of the α mitotype in different tissues of the *D. paulistorum* O11 line, we sequenced the O11 genome using Oxford Nanopore technology (ONT). The assembly of the nuclear genome of O11 was 231 Mb (111 contigs, N50 = 18.6 Mb) (**Table S3**), with a BUSCO completeness of 99% (diptera_odb, **Table S4**). Using the mitochondrial genome from the O11 line (β mitotype, Baião et al., 2023), we searched the nuclear assembly and found 12 mt-genome copies clustered at the end of one contig (ctg000940). From here on, we refer to this cluster as the *Dpau* Mega-NUMT, given its similarity to the theoretical Mega-NUMT concept. Further analysis of the 12 NUMT copies revealed a conserved synteny of the mitochondrial genes across most copies (**Fig. 3**). Four of the NUMT copies have lost genes compared to the mt-genome, and in all NUMT copies, many of the genes have suffered frameshift mutations (**Fig. 3**, **Table S5**). When comparing the gene length to the fraction of intact gene copies per protein-coding gene, we found a significant inverse correlation, suggesting that the mutation process is random (**Fig. S2**). However, given the complexity of the Mega-NUMT, it is possible that the sequence of each gene copy in the assembly is not 100% accurate. Nevertheless, we detect intact copies of the 13 different mitochondrial protein coding genes in the Mega-NUMT, albeit with the mitochondrial genetic code, and thus all contain stop codons when translated with the nuclear code (**Table S5**). We also detect copies in which insertions and deletions have caused frameshifts, which we refer to as pseudogenes **(Fig. 3, Table S5)**.

**Figure 3.**
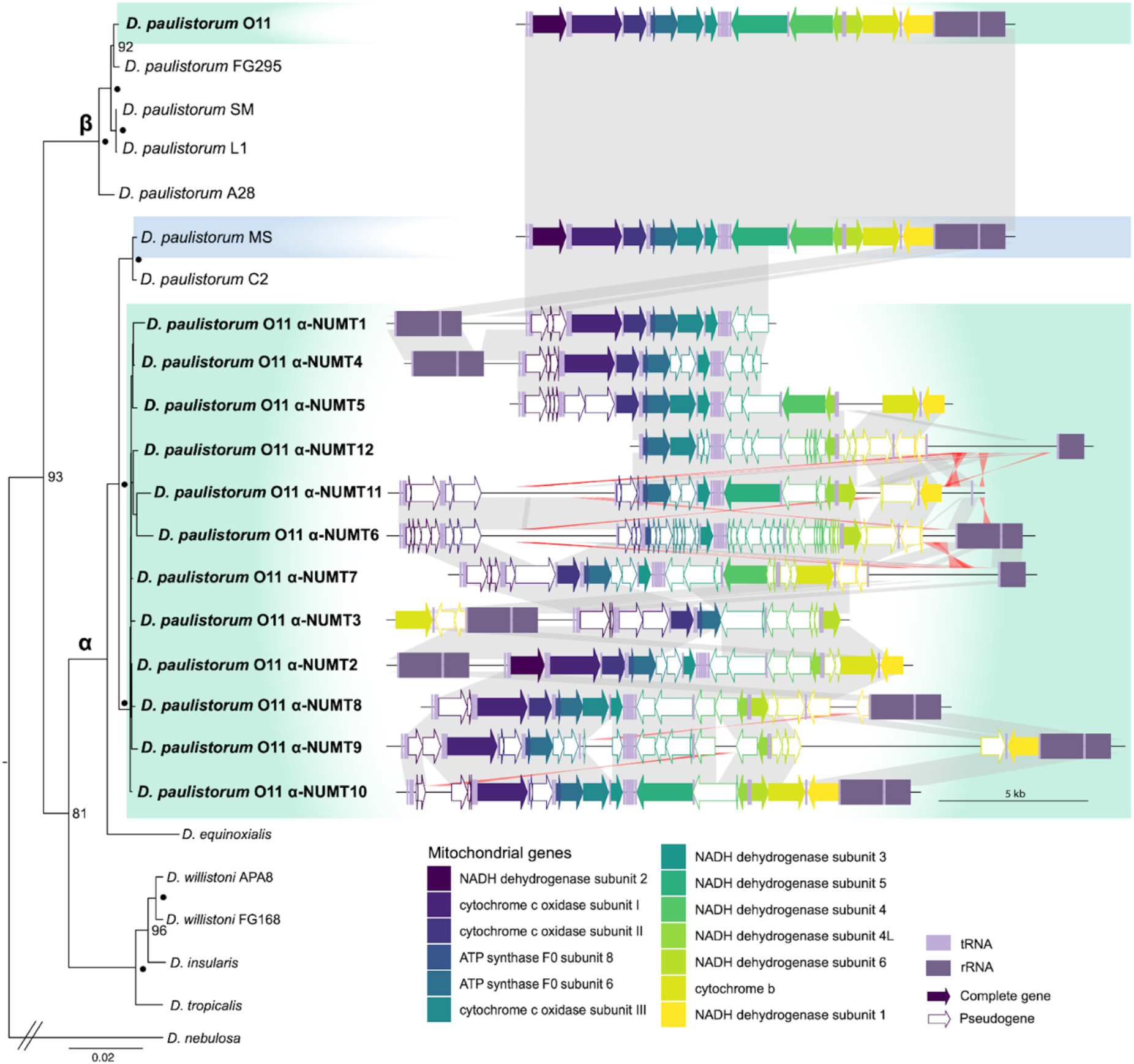
Phylogenetic position and synteny of NUMT copies of *D. paulistorum* O11 and closely related mitochondrial genomes. Maximum likelihood phylogeny based on the concatenated set of 13 genes and pseudogenes of the NUMT copies found in the Mega-NUMT of *D. paulistorum* O11, together with the mitochondrial genes in *D. paulistorum* and closely related species. Black dots indicate nodes with 100 bootstrap supports. The α and β mitotype clades are annotated on their respective nodes in the tree. Mitochondrial genomes of *D. paulistorum* O11 (green) and *D. paulistorum* MS (blue) were used as references for the β and α mitotypes. The NUMT copies of *D. paulistorum* O11 are highlighted in green to aid the visualization of the paraphyletic placement of *D. paulistorum* O11 in both β (as mitochondrial genome) and α mitotype lineages (as NUMTs). To the right of the tree, we show a representation of the synteny between the α-NUMT copies of *D. paulistorum* O11, the β mitotype mitochondrial genome of *D. paulistorum* O11 and the α mitotype of *D. paulistorum* MS. The grey synteny ribbons between sequences represent homologous regions in the same direction, while red ribbons represent inversions. The 13 mitochondrial protein-coding genes are shown as colored arrows. Genes with an intact open reading frame based on the mitochondrial genetic code are represented with filled arrows, while genes that have suffered frameshift mutations are represented as a white arrow with a colored outline, named pseudogenes. rRNA and tRNA genes are represented as rectangular blocks in two shades of pastel purple (rRNA, dark; tRNA, light).

A phylogeny based on the complete set of genes and pseudogenes in the *Dpau* Mega-NUMT, together with the published mitochondrial genomes of *D. paulistorum* spp. and other willistoni group species (Baião et al., 2023), shows that all individual NUMT copies in the Mega-NUMT are found in one clade, which is sister to the α mitotype of the *D. paulistorum* C2 and MS lines (**Fig. 3**) belonging to the CA and AB semispecies, respectively (**Table 1, Table S6**). Hence, we refer to these NUMT copies as α-NUMTs. Importantly, though, the separation of the α mitotype and α-NUMTs into different clades, and the monophyly within these clades, suggests that none of the α-NUMT copies in the *Dpau* Mega-NUMT are recent insertions from an extant α mt-genome. This is further supported by the fact that the O11 line carries the β mitotype, as does the majority of *D. paulistorum* semispecies and lines.

**Table 1.** α-NUMT abundance and prevalence in *D. paulistorum*. lines based on *CO1* RFLP-PCR from adult flies. Abundance is given in percentage of α-NUMTs compared to the mitochondrial copies per fly (**Fig S1**) and is presented separately for females (left) and males (right). Prevalence of the Mega-NUMT among tested individuals is presented both as a percentage and in absolute numbers (in parentheses). Abbreviations: Spp: Semispecies, AB: Andean-Brazilian, AM: Amazonian, CA: Centro-American, IN: Interior, OR: Orinocan, TR: Transitional, Coll: Collection year, ND: Not determined. Further details about the fly lines and genotyping are provided in **Table S6** and **Fig. S1**.

| Line | Coll. | Spp. | Mitotype | α-NUMT detected | α-NUMT abundance (%)<sup>#</sup> | α-NUMT prevalence |
| --- | --- | --- | --- | --- | --- | --- |
| O11 | 1957 | OR | $\beta$ | Yes | 30.7±3.7/36.8±3.6 | 100% (39/39) |
| FG103 | 2014 | OR | $\beta$ | Yes | ND<sup>§</sup> | 0% (0/40)<sup>§</sup> |
| FG111 | 2014 | OR | $\beta$ | Yes | 10.32±4.0/11.8±3.3 | 55% (22/40) |
| FG295 | 2014 | OR | $\beta$ | - | 0 | 0% (0/40) |
| TP37 | 2009 | OR | $\beta$ | - | 0 | 0% (0/20) |
| POA1 | 2003 | OR | $\beta$ | - | 0 | 0% (0/20) |
| RP | 1995 | OR | $\beta$ | - | 0 | 0% (0/20) |
| A28 | 1952 | AM | $\beta$ | - | 0 | 0% (0/40) |
| FG16 | 2018 | AM | $\beta$ | Yes | 57.2±8.6/66.5±7.2 | 55% (22/40) |
| FG18 | 2018 | AM | $\beta$ | - | 0 | 0% (0/40) |
| FG572 | 2015 | AM | $\beta$ | Yes | 45.3±8.3/63.2±6.5 | 83% (44/50) |
| MS* | 1962 | AB | $\alpha$ | Yes | 40.3±3.8/54.9±3.7 | 100% (20/20) |
| Yellow* | 1960 | AB | $\alpha$ | Yes | 20.4±4.7/27.4±4.3 | 100% (40/40) |
| C2 | 1954 | CA | $\alpha$ | - | 0 | 0% (0/20) |
| White | 1960 | CA | $\alpha$ | - | 0 | 0% (0/20) |
| L1 | 1958 | IN | $\beta$ | - | 0 | 0% (0/20) |
| SM | 1956 | TR | $\beta$ | - | 0 | 0% (0/20) |

<sup>#</sup> Abundance of the α-NUMT is given in percentage of females and males determined by *COI* RFLP-PCR.
<sup>§</sup>The line FG103 presented α-NUMT in 2014/15 (determined by Sanger and Illumina sequencing) but has lost it afterwards (verified by *COI* RFLP-PCR and Sanger sequencing in 2018/19).
<sup>\*</sup>To distinguish $\alpha$ NUMTs from the α-mitotype of AB semispecies, the α-diagnostic *EcoRV* site of *COI* was used (Fig. S1).

Further, we analyzed the genomic region where the *Dpau* Mega-NUMT is located. The Mega-NUMT is directly flanked by an AT-rich 36 bp satellite sequence (**Fig. 4** and **Fig. S3**) and the gene *FOXO* encoding the Forkhead box O transcription factor (Jünger et al., 2003), located in Muller element E on chromosome 3, as deduced by comparison to *D. melanogaster*. The same 36 bp satellite DNA is also found inside the Mega-NUMT, downstream of α-NUMT 3 (**Fig. 4** and **Fig. S3**). The positions of *FOXO* and the Mega-NUMT next to each other were also confirmed by DNA-hybridizations on polytene chromosomes of larval salivary glands of O11, and the bright polytenized signal of the Mega-NUMT (**Fig. S4**) implies its euchromatic state since it is polytenized.

**Figure 4.**
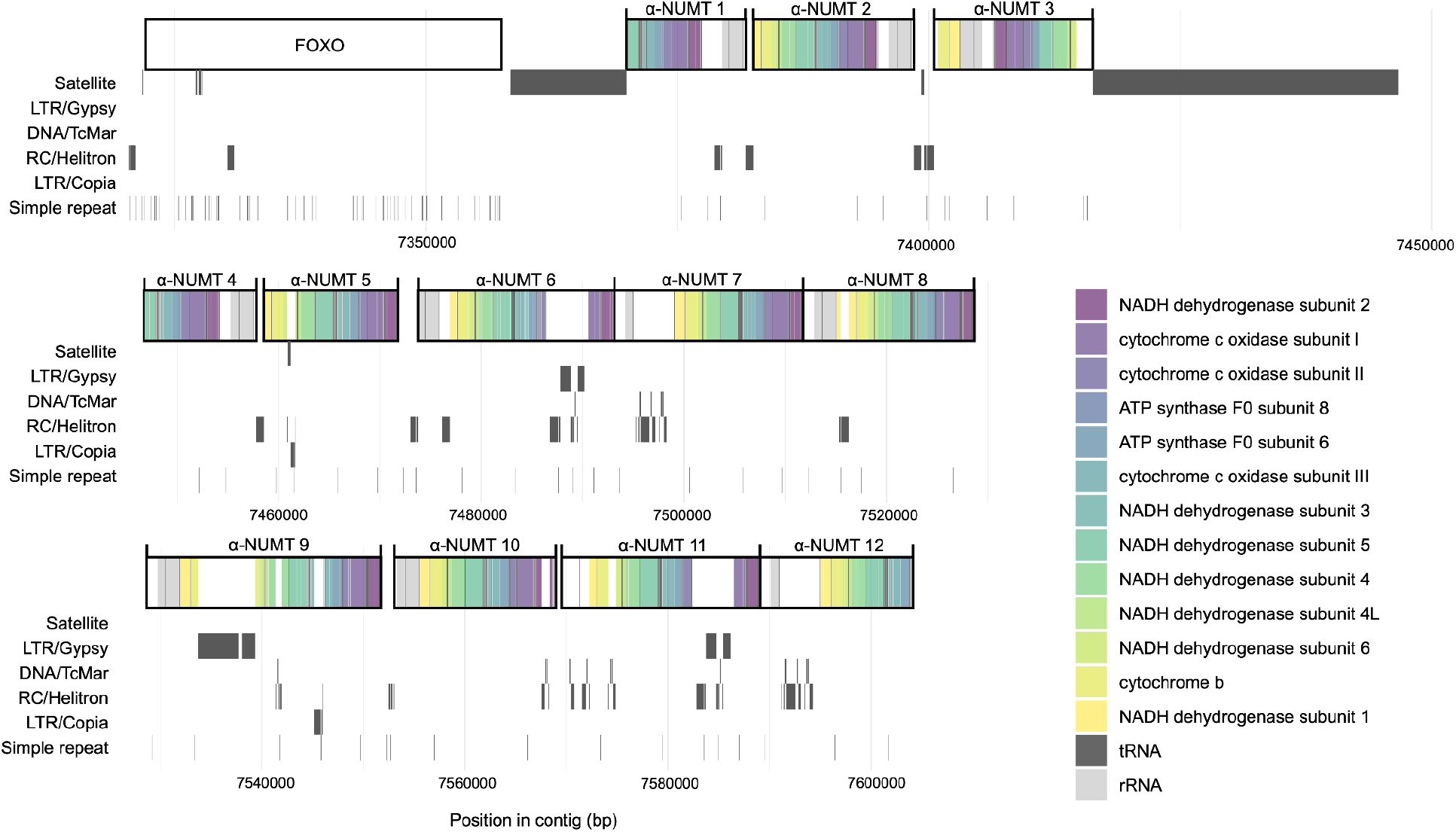
Graphic representation of the end of contig000940 in the assembly of *D. paulistorum* O11. The region is divided into three rows to facilitate visualization. The position of the nuclear gene *FOXO* and each α-NUMT is represented with a white box, which also contains colored lines that represent the position of each protein-coding and RNA gene. The positions of the satellite and repeat elements are shown below in the order: The AT-rich 36-bp satellite, *Gypsy*, *TcMariner*, *Helitron*, *Copia*-like transposons, and simple repeats.

All α-NUMT copies except α-NUMT 3 have the same orientation in the *Dpau* Mega-NUMT (**Fig. 4**). The sequence of α-NUMT 3 also contains a rearrangement not seen in any of the other copies of the contig (**Fig. 3** and **4**). Additionally, the 12 α-NUMT copies in the O11 nuclear genome assembly are surrounded by repeat elements, and in some cases, insertions of transposable elements are present within an α-NUMT sequence (**Fig. 4**). Specifically, we found remnants of the LTR retrotransposon *Gypsy* within the α-NUMT copies 6, 9, and 11 and of the LTR element *Copia* in α-NUMT copies 5 and 9 (**Fig. 4**). In addition, we found short regions with sequence similarity to the DNA transposon Tc Mariner in six of the α-NUMTs and to the rolling-circle DNA transposable element, *Helitron*, in nine of the α-NUMTs, as well as between α-NUMT copies when a non-NUMT sequence was found between them (**Fig. 4**).

### Number of α-NUMT copies in the genome

Since the discriminant qPCR results indicated that more than 12 α-NUMT copies could be present, we investigated the coverage of ONT reads in the Mega-NUMT region of O11 to assess the true number of α-NUMT copies. When mapping the reads to the assembly, we also included the β mt-genome from Baião et al., (2023) and the genome of the bacterial symbiont *Wolbachia* of O11 from Papachristos et al., (2025) to prevent mapping of reads from the real mitochondrion or *Wolbachia* to the Mega-NUMT. Additionally, to be as stringent as possible in our assessment, only one mapping per read was allowed. The read coverage over the *Dpau* Mega-NUMT shows a clear discordance with the average coverage across the genome (**Fig. 5**). The *Dpau* Mega-NUMT has an average coverage of 402±159, while the genome assembly as a whole has an average coverage of 80.8±77.4 (**Fig. 5**). Using these numbers, we thus infer that the genome contains around 60 (402/80*12 copies) whole mt-genome α-NUMTs (**Fig. 5**). We note that the coverage over the different α-NUMT copies varies substantially. Especially high coverage was found on α-NUMT 8, with 17 times higher coverage than the genome average, followed by α-NUMTs 7, 9, and 10, with ten, seven-, and six-times higher coverage than the genome average, respectively. These results indicate that the unassembled α-NUMT copies resemble α-NUMT copies 7-10 the most. Specifically, we note that both α-NUMT 7 and 8 contain the full mt-genome and have no intervening sequence, suggesting that many of the unresolved copies are also full mt-genomes with no intervening sequences (**Figs. 4** and **5**).

**Figure 5.**
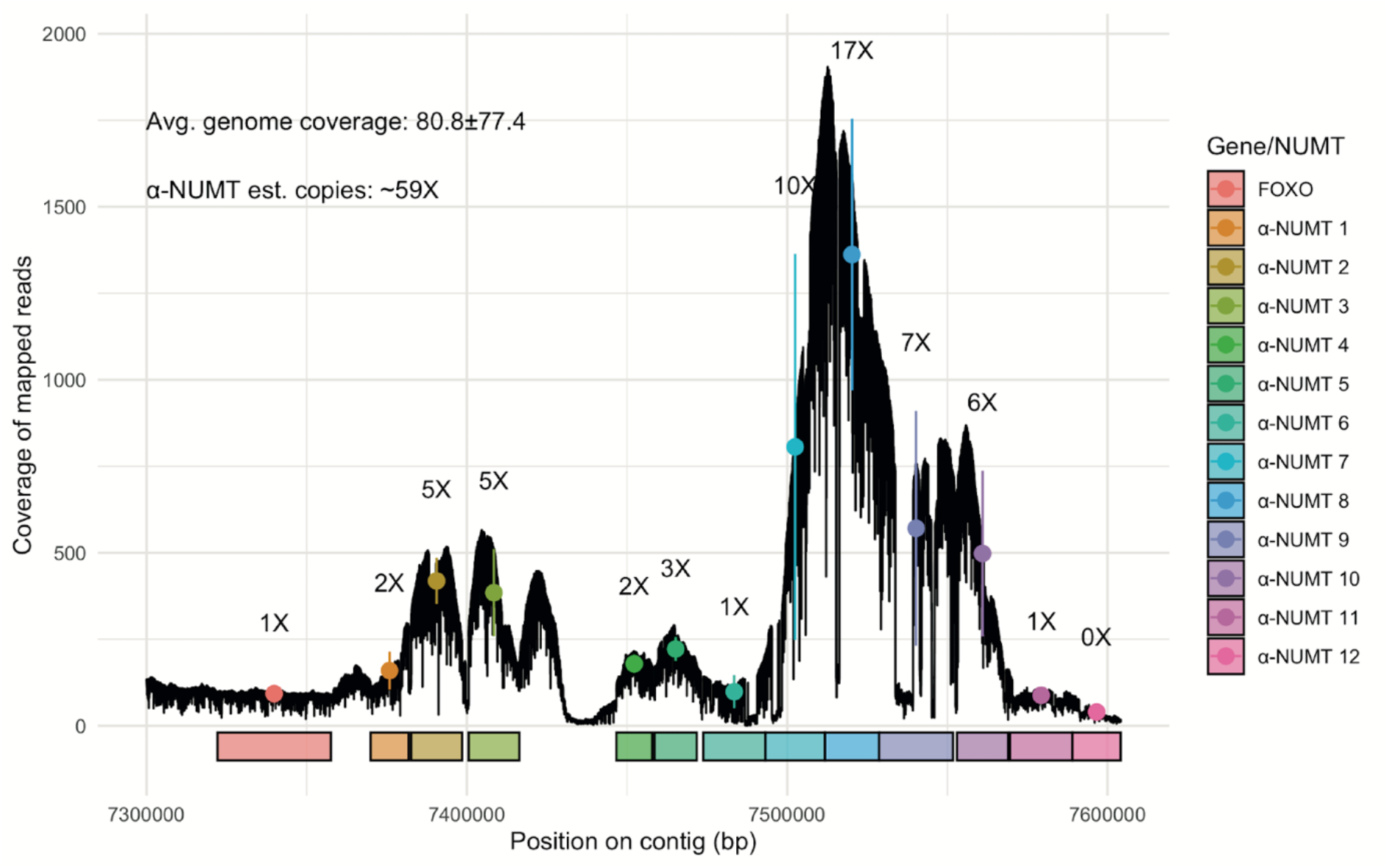
Coverage quantification of α-NUMTs in the nuclear genome of *D. paulistorum* O11. The x-axis represents the end region of contig000940 of the genome assembly of *D. paulistorum* O11, which includes the nuclear gene *FOXO*, and the 12 assembled α-NUMTs. Coverage was obtained from mapping Nanopore reads to the genome assembly. The average and standard deviation of the coverage over each α-NUMT and the *FOXO* gene are plotted in the same color as used for the box depicting each feature. The total number of α-NUMT copies in the Mega-NUMT is estimated from the average over all the assembled α-NUMT copies in relation to the average coverage of the whole genome assembly.

We note that the 60 α-NUMT copies estimated from long read mapping are more than the ca. 30 we estimated by our qPCR results. As α-NUMT 3 lacks *CO3* completely, only 11 of the 12 copies in our assembly have a binding site for the primer CO3_SNP_1 used in qPCR for determining the copy numbers (**Fig. 2**, **Table S1**). Hence, although we do not know exactly what the remaining copies look like, this indicates that the discrepancy in total copy numbers of the α-NUMT between the coverage and qPCR results might be explained by a lack of primer binding in some of the α-NUMT copies, either due to the primer binding site being absent or containing mutations. However, given that the coverage is relatively noisy, we cannot fully determine the number of α-NUMT copies in the O11 genome. Instead, we estimate that there are between 30-60 nearly complete α-NUMT copies in the *Dpau* Mega-NUMT.

### Mega-NUMT visualization on *D. paulistorum* chromosome

To validate the genomic findings of the Mega-NUMT localization on the 3^rd^ chromosome of *D. paulistorum* O11, we performed DNA fluorescent *in situ* hybridization (FISH) on mitotic chromosomes of larval brains using mitochondria-specific probes and a probe for the Mega-NUMT-linked satellite (see **Fig. 4** and the Materials and Methods section). In O11, we observed two satellite signals (red) on the 3^rd^ chromosome—one in the (peri)centromeric region and a second in the subtelomeric region—as well as a Mega-NUMT signal (green) adjacent to the subtelomeric satellite signal (**Fig. 6A**), in agreement with the assembly data (**Fig. 4**). In contrast, the *D. paulistorum* line FG295 (**Table S6**), which also belongs to the OR semispecies (**Table 1**), shows only a single satellite signal in the (peri)centromeric region of the third chromosome, but no Mega-NUMT signal (**Figure 6B**).

**Figure 6.**
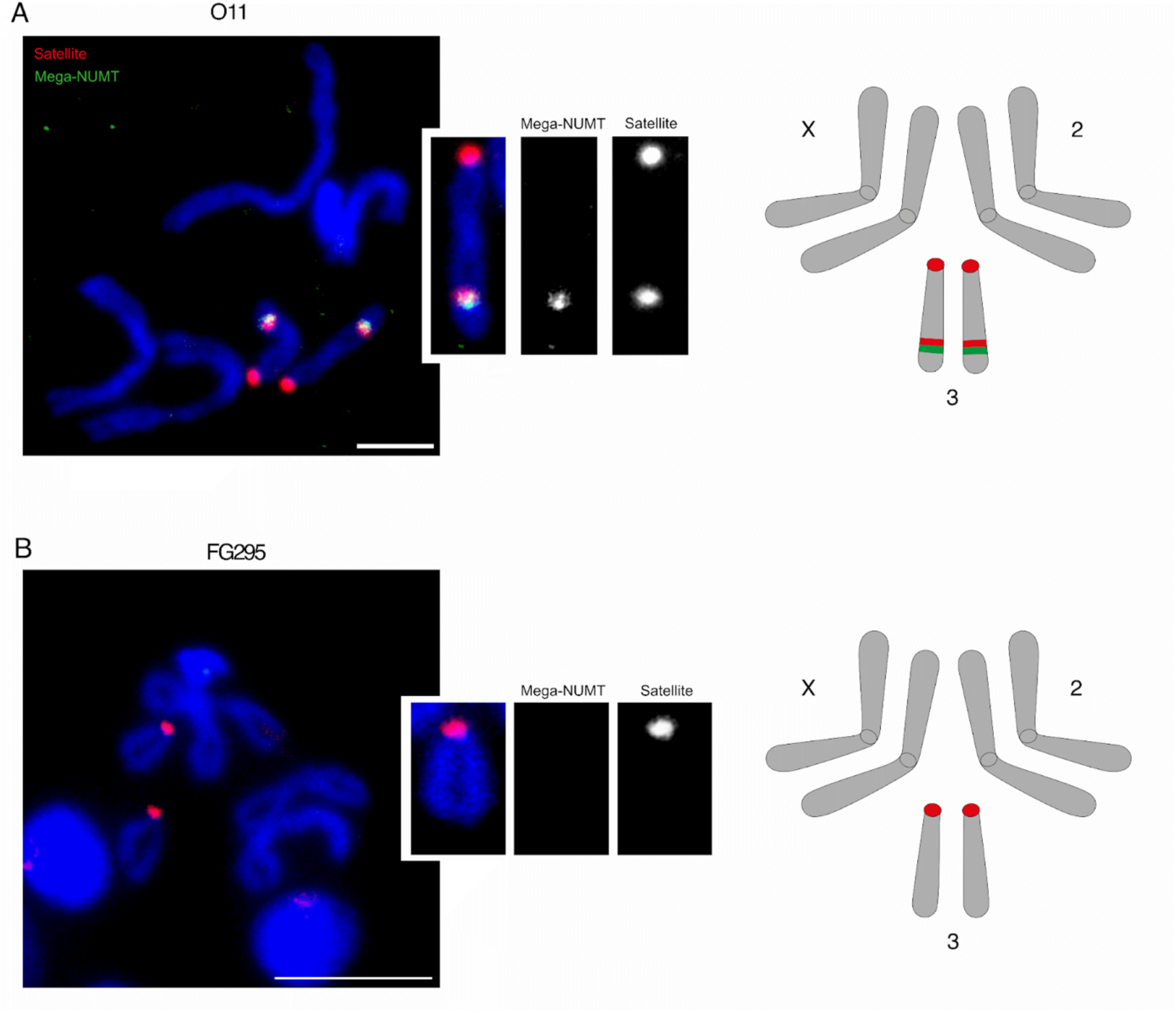
DNA-FISH on mitotic chromosomes from larval brains of the *D. paulistorum* O11 and FG295 lines. Probes targeting the Mega-NUMT (green) and the Mega-NUMT–linked satellite (red) were used. The scale bar represents 5 μm. The inset highlights the third chromosome, showing two satellite signals and one Mega-NUMT signal in O11 (**A**), but only a single satellite signal in FG295 (**B**). The schematic on the right summarizes the localization of the satellite (red) and the Mega-NUMT (green) in each strain.

Additionally, to follow Mega-NUMT dynamics in interphase and dividing cells, we investigated the localization of the *Dpau* Mega-NUMT in early embryonic mitosis by mt-DNA FISH (**Fig. S5**). Contrary to larval brains that were generated by squash fixation and thereby washing off the cytoplasm (**Fig. 6**), in our whole mount confocal mt-DNA-FISH assays on O11 embryos, we observed staining of mitochondria in the cytoplasm and the chromosome-associated Mega-NUMT signal as two dots, with some cases of a fused, larger dot (arrowheads and arrow, respectively, **Fig. S5A-D, F**). Each dot consists of two parts with a small bridge in between, suggesting somatic pairing (inset of **Fig. S5B**). Based on our results on metaphase chromosomes from 3^rd^ instar larvae (**Fig. 6**), we conclude that each dot represents a Mega-NUMT signal located on one of the duplicated arms of chromosome 3, with two smaller parts representing sister chromatids. At the anaphase stage, we could clearly observe the separation of sister chromatid signals, creating a 4-dot pattern (**Fig. S5E**). Altogether, systematic DNA-FISH at different stages of early embryo mitosis reconfirmed the presence of only one Mega-NUMT signal on chromosome 3 of the O11 line.

### Prevalence of the Mega-NUMT across *D. paulistorum* semispecies and populations

To test if the *Dpau* Mega-NUMT is restricted to only the O11 line, we analyzed α-NUMT prevalence across different *D. paulistorum* lines. We used RFLP-PCR from adult flies to assess the prevalence and abundance of α-NUMTs (**Table 1**). In addition to O11, another OR line, FG111, presented α-NUMT, but it appeared not to be fixed in the population, as it was only found in 10 out of 40 flies that originated from an isofemale (**Table 1**). In yet another OR line, FG103, the Mega-NUMT was lost between the first test (2014-15) and the last one (2018) (**Table 1**). Two AM lines, FG16 and FG572, also presented a non-fixed Mega-NUMT (**Table 1**), while two lines from the AB semispecies, MS and Yellow, both carried the Mega-NUMT in all tested flies (**Table 1**). To sum up, we found the Mega-NUMT in several OR lines, as well as in the AM and AB semispecies, but not in the few representatives of the other three semispecies of *D. paulistorum* (CA, IN and TR). Possibly, this is caused by a sampling bias, since none of these three semispecies were found in our collections from French Guiana between 2014 and 2018 (**Table 1** and **Table S7**). In addition to O11, we only found fixation of the *Dpau* Mega-NUMT in the old long-term stocks of MS and Yellow that were maintained at small population sizes in the laboratory since the 1960s or earlier. Our finding that the Mega-NUMT is not fixed in isofemale lines from our recent collections in French Guiana suggests that it is frequently hemizygous in wild populations.

To verify homozygosity of the Mega-NUMT in the O11 line, we mated FG295 females that do not carry the Mega-NUMT (**Table 1**) to O11 males and determined that all F1 progeny inherited the Mega-NUMT (**Fig. S6**).

To evaluate the prevalence of the *Dpau* Mega-NUMT in nature, we screened a total of 80 *D. paulistorum* spp. isofemale lines at generation F1 post-collection that were sampled between 2014 and 2024 from five different locations in French Guiana (**Fig. 7**; **Table S7**).

**Fig. 7.**
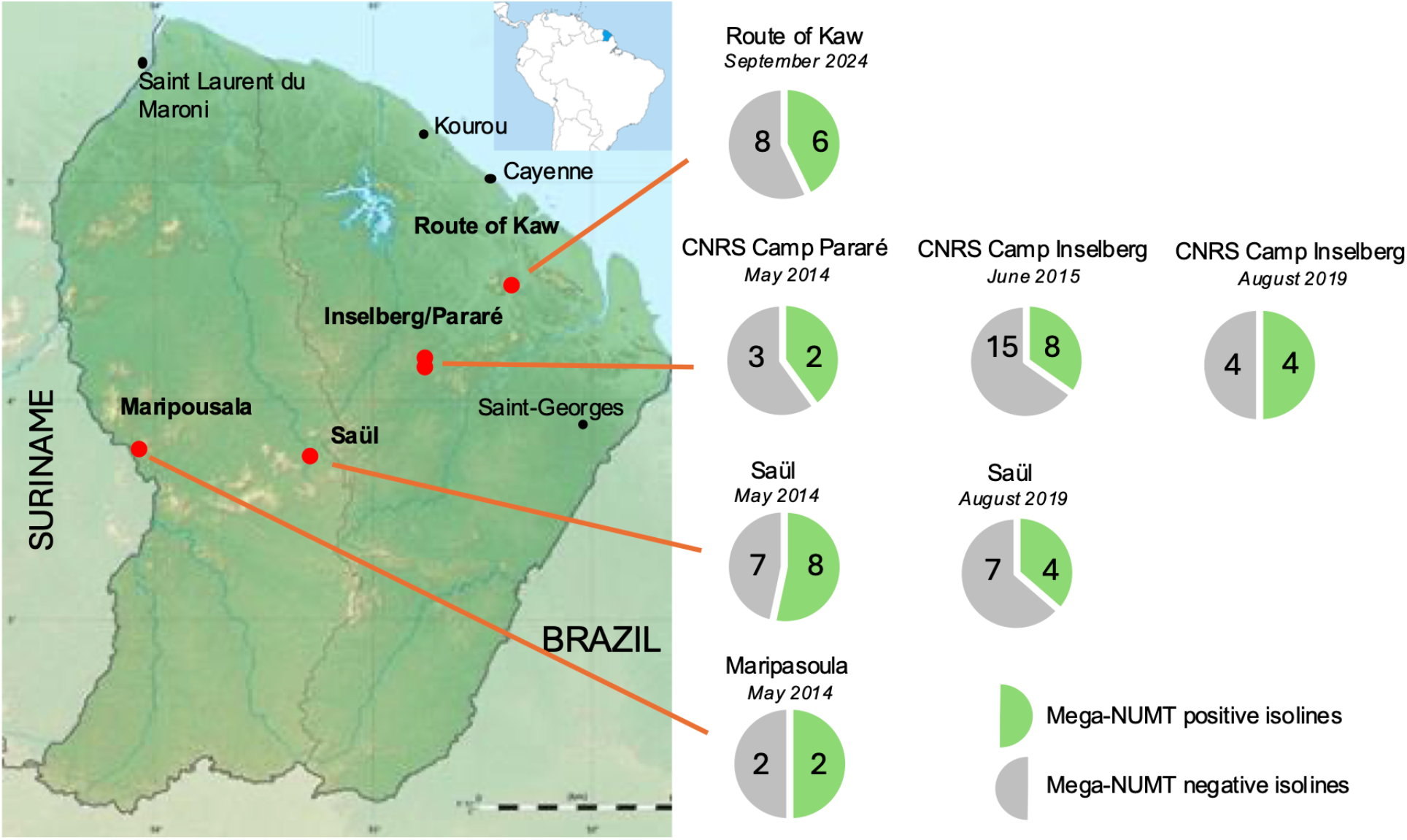
Natural prevalence of Mega-NUMTs in *D. paulistorum* spp. populations in French Guiana. All 80 fly samples were collected between 2014 and 2024 at five different locations (red dots, and **Table S7**). The individual pie charts are in reference to locality and collection date and show the actual number of Mega-NUMT-positive (green) and negative (gray) specimens. Species designation and Mega-NUMT-typing per established *D. paulistorum* spp isoline (n=80) was performed on multi-fly DNA at generation F1 post-collection by *CO1* PCR and direct sequencing. The presence of robust double peaks in their chromatograms at α-NUMT-diagnostic positions was used in combination with cloning and sequencing.

Sampled flies were established as isofemale lines and barcoded at F1/F2 post-collection as members of the *D. paulistorum* species complex by *CO1* PCR followed by direct Sanger sequencing and/or cloning of *CO1.* All *D. paulistorum* flies sampled in French Guiana are belonging either to the OR or AM semispecies, but never the Andean Brazilian (AB) one with the α1/α2 mitotype, (**Table S7**). Similar to O11 (**Fig 1A**), the presence of robust double peaks in the chromatograms from direct sequencing was used for Mega-NUMT typing, suggesting that 34 of the 80 isofemale lines harbor α-NUMTs (**Table S7**). The high prevalence of α-NUMT of 43% in our recent collections from *D. paulistorum* populations, together with its presence in at least three of the six semispecies (**Table 1**), strongly suggests their ancestral evolutionary state in this neotropical species complex and its stable maintenance both in nature and over long evolutionary time.

Finally, to test the stability of the *Dpau* Mega-NUMT under long-term lab conditions, we randomly selected 31 isolines, collected in French Guiana between 2014 and 2015, for *CO1* resequencing (**Table S8**). From these, more than half (16/31) were originally typed Mega-NUMT positive (51.6%), whereas after more than 10 years, only 7 out of the 16 lines were still positive, suggesting that more than half of the Mega-NUMTs were lost. The significant drop in MN prevalence in our stock over lab culturing from 51.6% (16/31) to 22.6% (7/31) suggests that the Mega-NUMT may drift to either fixation or loss under laboratory conditions, where selection is reduced or different than in nature.

## Discussion

In animal species, mtDNA is normally transmitted strictly uniparentally from mother to progeny (Ladoukakis & Zouros, 2017), with rare exceptions (Breton et al., 2007). However, reports of mitochondrial heteroplasmy and biparental transmission in humans challenged this view (Schwartz & Vissing, 2002; Luo et al., 2018) and prompted skepticism by the scientific community (McWilliams & Suomalainen, 2019; Salas et al., 2020). Consequently, the theoretical Mega-NUMT concept, proposing nuclear integration of multiple copies of whole mitochondrial genomes, was coined as an alternative explanation for such unorthodox findings (Balciuniene & Balciunas, 2019). Although this concept has been verified in humans (Wei et al., 2020; Lutz-Bonengel et al., 2021; Pagnamenta et al., 2021; Wei et al., 2022), similar phenomena have so far only been described in a limited number of animal systems where only partial regions of the mt-genome were found at multicopy (Lopez et al., 1994; Patterson et al., 2023; Biró et al., 2024), but not in Drosophila (Rogers & Griffiths-Jones, 2012), despite a relatively high prevalence of NUMTs in insects (Hebert et al., 2023). On the other hand, nuclear insertions of whole bacterial genomes larger than 1 Mb, like the α-proteobacterium *Wolbachia*, have been repeatedly detected in invertebrate genomes, including insects and nematodes (Kondo et al., 2002; Hotopp et al., 2007; Nikoh et al., 2008; Klasson et al., 2014; and recently reviewed in Keeling, 2024).

### The Mega-NUMT of *D. paulistorum* spp

In this study, we found that multiple whole mitochondrial genomes are integrated in the nuclear genome of the O11 line from the *D. paulistorum* species complex, designated α-NUMTs because of their close phylogenetic relationship with the cytoplasmic α-mitotype of the Centro-American and Andean-Brazilian semispecies of *D. paulistorum* (Baião et al., 2023). These α-NUMTs localize in a nuclear cluster in the Müller element E on chromosome 3 with up to 60 nearly complete 15kb mitochondrial genomes, potentially with a total size of 900 kb. Given the agreement between this finding and the Mega-NUMT concept, we named this cluster the “*Dpau* Mega-NUMT”. Similar to earlier NUMT reports (Behura, 2007; Tsuji et al., 2012; Dayama et al., 2014; Schiavo et al., 2017; Wang et al., 2020; Biró et al., 2024), the *Dpau* Mega-NUMT of the O11 line is located in a genomic region containing other repetitive elements, *i.e.,* a chromosome 3-specific satellite and several different TEs (**Fig. 6A, B**), which suggests it might have served as a “safe haven” for such sequences (Werren, 2011). Additionally, contrary to currently described human Mega-NUMTs with prevalence below 0.5% (Wei et al., 2020; Bai et al., 2021; Lutz-Bonengel et al., 2021), the *Dpau* Mega-NUMT is found at high prevalence, around 40% in long-term laboratory lines (3/8; **Table 1**) and 42.5% (34/80) in *D. paulistorum* spp. flies collected between 2014 and 2024 at different locations in French Guiana (**Fig. 7**; **Table S7** and **Table S8**). Furthermore, the *Dpau* Mega-NUMT is present in members of at least three of the six semispecies of *D. paulistorum*, either in homo- or hemizygosity (**Table 1**). Finally, the α-NUMTs form a separate monophyletic clade to the exclusion of the current α mitotype (**Fig. 3**), indicating that the insertion did not involve any of the extant mitotypes but possibly a more ancestral one. Therefore, we proposed that the *Dpau* Mega-NUMT might have been inserted in the nuclear genome before or just after the semispecies radiation, estimated to have occurred between 1.2-2.1 MYs ago (Zanini et al., 2018; Kim et al., 2022).

### Integration and chromosomal position

The overall well-conserved head-to-tail structure of the *D. paulistorum* α-NUMT cluster may suggest that the *Dpau* Mega-NUMT could have integrated already as a concatemer of multiple mt-genomes and not as a single unit. As earlier proposed by Balciuniene & Balciunas, 2019, the mechanism of generating Mega-NUMTs of complete mt-genomes may be explained by replication via a rolling circle intermediate. Such mt-rolling circles have, for example, been reported in yeast, nematodes, and humans (Pohjoismäki et al., 2009; Ling et al., 2016; Ling & Yoshida, 2020), but not in Drosophila until now. The switching between conventional circular to rolling circle replication might be quite feasible because key components of the mitochondrial transcription and replication apparatus are derived from the T-odd lineage of bacteriophage rather than from an α-Proteobacterium, as the endosymbiont hypothesis would predict (Filée et al., 2002; Shutt & Gray, 2006). In addition, rolling circle replication (RCR) has been described in Drosophila for *histone* and the *Stellate* and *Suppressor of Stellate (Su(Ste))* genes (Cohen et al., 2005; Cohen & Segal, 2009). Moreover, *Helitron* transposons like *DINE-1* elements that can reach extremely high copy numbers in *D. melanogaster* (Thomas et al., 2014) also propagate via RCR (reviewed in Barro-Trastoy & Köhler, 2024).

The nuclear integration site of the mt-concatemer in the chromosome 3 of *D. paulistorum* spp. is interesting, since it is a fusion between Muller elements E and F, orthologous to chromosome arm 3R and chromosome 4 of the melanogaster group, respectively (Muller, 1940; Spassky & Dobzhansky, 1950). The location of the *Dpau* Mega-NUMT in the O11 line next to the satellite DNA (**Fig 6A**), also present in the acrocenter of the fusion autosome E/F (**Fig. 6B**), potentially suggests a chromosomal inversion event splitting the satellite in two, but not affecting the upstream gene *FOXO*. Such paracentric inversions are quite commonly found in the *D. paulistorum* species complex (Kastritsis, 1967) and can even play adaptive roles (Krimbas & Powell, 1992), like for the successful recent colonization of urban environments (Valiati & Valente, 1997). In addition, inversions are enriched on chromosome 3 and often maintained in heterozygosity, potentially under balancing selection (Gromko & Richmond, 1978). These earlier findings suggest that chromosome 3 of *D. paulistorum* spp. may be an evolutionary hotspot for chromosomal rearrangements like inversions and translocations, but also mitochondrial-derived insertions like the *Dpau* Mega-NUMT as reported here.

### Why is the *Dpau* Mega-NUMT maintained at high prevalence and over evolutionary time?

Contrary to currently described human Mega-NUMTs, which are extremely rare in prevalence, the *Dpau* Mega-NUMT appears to be evolutionarily persistent since it is present in lines from at least three different *D. paulistorum* semispecies, *i.e*., Amazonian, Andean Brazilian and Orinocan. Additionally, it is present at a relatively high prevalence of potentially more than 40% in natural populations (**Figure 7**; **Table S7**). Even so, the fact that the Mega-NUMT is not fixed in *D. paulistorum* except for old laboratory stocks (**Fig. 1**; **Table 1**), and that natural populations are mainly hemizygous, may suggest it is maintained under some kind of balancing selection rather than having an essential function. Moreover, selective constraints may differ in nature from laboratory conditions over 10 years (exposed to drift and inbreeding for more than 200 fly generations) as more than 50% of the initially FG lines collected between 2014 and 2015 had lost their Mega-NUMTs, at least they were no longer tracible by direct Sanger sequencing (**Table S8**). Follow up studies at the population level will be needed to further elucidate their nature and evolution.

Although almost half of the mitochondrial genes in Mega-NUMT present in the 12 resolved α-NUMT copies in our assembly (**Table S5**) still appear to encode intact ORFs (albeit with the mt genetic code), we believe that they are unlikely to be under selection for having the same function as when they were present in the mt-genome. Mainly because the mt genetic code will cause problems during translation, but also since other mechanisms differ between the mitochondria and nucleus regarding gene expression. Additionally, although we can’t fully determine whether some of them might be due to sequencing errors, it appears as if the pseudogenizing mutations occurred randomly throughout the genes (**Fig. S2**), suggesting neutrality. However, we can’t exclude that some of the DNA in the Mega-NUMT might be transcribed and serve yet undetermined functions. Potentially, the TEs within and between α-NUMT units, especially remnants of LTR retrotransposons like *Copia* and *Gypsy* (**Fig. 4**), may contribute to expression, as LTRs are well known to harbor cell- and tissue-specific *cis*-regulatory elements that affect expression of neighboring host genes (McDonald et al., 1997). Alternatively, TEs can also create *de novo* regulatory regions acting *in trans*, so-called piRNA clusters (recently reviewed in Pritam and Signor, 2025), which, if expressed in the antisense orientation of the transposon, can serve as efficient epigenetic silencers of homologous TEs via the piRNA pathway (Brennecke et al., 2008). Indeed, *de novo* creation of such a repressive piRNA cluster has recently been demonstrated in *D. simulans* (Rafanel et al., 2025). However, currently, these are purely speculations that need to be tested further.

Another possibility is that the Mega-NUMT provides a structural function. Other studies have seen that NUMTs are located close to centromeric or telomeric regions (Puertas & González-Sánchez, 2020), suggesting that they might be involved in their function, as both are normally consisting of repetitive sequences that can be highly species-specific. However, the *Dpau* Mega-NUMT of O11 is not found in either the centromeric or telomeric chromosomal region of chromosome 3, according to our FISH. Even so, a yet undetermined structural role for the *Dpau* Mega-NUMT is still possible.

PacBio HiFi and Hi-C data from additional genomes will be necessary to elucidate the evolution and structure of the Mega-NUMT, and further experiments are also needed to determine potential functions and fitness benefits of the Mega-NUMT in the *D. paulistorum* species complex.

## Conclusions

To our knowledge, this is the first verified existence and detailed dissection of a Mega-NUMT outside cats and humans. Moreover, we show that Mega-NUMTs can persist at high prevalence in nature and over relatively long time periods. Although we have not tested this hypothesis more explicitly, this pattern suggests some kind of balancing selection. Our findings strengthen the importance of high-quality long-read sequencing technologies for deciphering complex repeat-rich genomic regions and mobile DNAs to deepen our understanding of genomic “dark matter”. Finally, it is expected that with the rapidly increasing number of high-quality genomes, the *Dpau* Mega-NUMTs will not remain a unicorn. Instead, similar Mega-NUMTs will very likely be found in further eukaryotic genomes.

## Data availability

The O11 genome assembly and the associated ONT reads are available in ENA with accession number PRJEB111780.

## Acknowledgements

We thank Tom Martin for performing the DNA extraction for Nanopore sequencing of *D. paulistorum* O11 and Catarina Gameiro for excellent technical support in fly line maintenance and their molecular-typing. ONT sequencing of O11 was performed by the Uppsala Genome Center (UGC) in Uppsala, Sweden. The facility is part of the National Genomics Infrastructure (NGI) Sweden and the Science for Life Laboratory. Some of the data handling was enabled by resources provided by the Swedish National Infrastructure for Computing (SNIC) at UPPMAX. UGC and UPPMAX are supported by the Swedish Research Council. This work was supported by the Swedish Research Council VR grant 2014–4353 to LK and by the Austrian Science Fund FWF grants P28255B22, FW613A0501 and FW613A0502 to WJM. We also thank the Nouragues research field station (managed by CNRS), which benefits from “Investissement d’Avenir” grants managed by Agence Nationale de la Recherhe (AnaEE France ANR-11-INBS-0001; Labex CEBA ANR-10-LABX-25-01) to WJM.

## Contributions

LK and WJM conceived and designed the study. MMN performed assemblies, annotations and phylogenetic analyses. EH performed qPCRs. JT and DIS performed line screening and RFLP analyses. AS and CC performed FISH. AH-V and WJM collected flies. LK performed bioinformatic analyses. MMN, WJM and LK wrote the paper with contributions from CC and AS. All authors read and approved the manuscript.

## Material and Methods

### Fly strains

All fly lines used in the present study are listed in **Table S6**, including their collection date and semispecies designation. All flies were kept at 25±1°C on Formula 4-24 Drosophila instant food (Carolina, USA) with a 12-hr light-dark cycle.

### Drosophila collections from French Guiana 2014-2024

Field samples of neotropical Drosophila specimens were collected in French Guiana at five different geographic locations between 2014 and 2024 (**Table S7**) by trapping live flies with banana/yeast baits in plastic PET bottles over a 48 hrs. time period. Taxonomic grouping - at least at the species-group level - was performed by eye under the stereo loupe by using FlyNap. Trapped inseminated single females were isolated into individual small single fly culture vials supplemented with Carolina Instant Fly-Food, shipped back to the Medical University of Vienna to establish consecutive isofemale lines in WM’s lab. Upon egg deposition and the emergence of F1 larvae, DNA was isolated from the G0 mother for single fly DNA extraction with the Gentra Puregene Tissue kit (Qiagen, Germany) and mitochondrial *CO1*-barcoding. PCR was performed with the primer pair CO1-F and CO1-univR (**Table S1**), followed by direct Sanger sequencing and cloning (for further details, see Madi-Ravazzi et al., 2021).

This study was conducted under ABS Permits No. ABSCH-IRCC-FR-245930-1 and ABSCH-IRCC-FR-257164-1, issued by the French Ministry of Ecological Transition on 25 Jun 2018 and 16 Sep 2021 respectively, in compliance with the Nagoya Protocol and French Biodiversity Law (2016).

### Single fly Southern blots

Single fly Southern blots were performed following the protocol by Junakovic, 2004. In brief, DNA was extracted from single individuals, digested with the restriction enzyme *Hin*dIII and processed for electrophoresis on a vertical 0.8% agarose gel. After vacuum-blotting onto a positively charged nylon membrane, fragments were hybridized with a P^32^-labelled mitochondrial *CO1* probe derived from a plasmid. Detection was performed by exposing the membranes to X-ray films at −80°C.

### Restriction Fragment Length Polymorphism-PCR (RFLP-PCR)

Total DNA was extracted from single or multiple flies using the Gentra Puregene Tissue kit (Qiagen, Germany). Flies were homogenized using a TissueLyser LT (Qiagen, Germany) at 50 Hz for 50 sec and DNA was consequently isolated following the manufacturer’s protocol. DNA concentration was determined with a NanoDrop OneC Spectrophotometer (Thermo Scientific, USA). RFLP-PCR for the mitochondrial Cytochrome c oxidase genes *CO1* and *CO2* (**Figure S1**) was set up in 20 µl reactions containing 1x Promega reaction buffer, 2.5 mM MgCl2, 0.5 μM of each primer (**Table S1**), 200 μM of each dNTP, and 0.025 U of GoTaq G2 DNA Polymerase. *CO1*/*CO2* reactions were performed using the following thermal profile: initial denaturation for 2 min at 95°C followed by 30 cycles consisting of 45 sec denaturation at 95°C, 45 sec annealing at the appropriate primer conditions T_m_°C (**Table S1**) and 30 sec extensions at 72°C. Final extension was carried out at 72°C for 10 min. PCR products were consequently used for RFLP (digest plus undigested control) using the restriction enzymes *Hin*dIII and *Eco*RV-HF for *CO1* (**Figure S1**). Reactions were performed in 20 µl volume containing 10 or 20 U of enzyme, 1x reaction buffer and 10 µl PCR product. After overnight incubation at 37°C, enzymes were inactivated by either heat (80°C for 20 min) or by adding EDTA-containing gel loading dye. 10 μl of undigested and digested samples were then run on a 1.5% agarose gel and post-stained with 10 mg/ml ethidium bromide. Imaging was performed using a Molecular Imager ChemiDocTM XRS Imaging System and the intensities of obtained PCR bands were quantified using Image Lab 5.2.1 software (Bio-Rad).

### Quantifying Mega-NUMT abundance within a fly

Undigested samples representing the sum of both mitotypes were set as a reference band with relative quantity 1. The relative levels of mitotypes were then quantified by summing up the two digested fragments plus the undigested fragment. Due to technical differences in band intensity, however, the sum of all fragments did not result in a relative quantity of 1. Hence, the actual sum value was set to 100% to determine the percentage of mitotype levels. Smaller fragments with a length up to approximately 100 bp were not detectable after ethidium bromide staining and we hence used the correction factor C*f* = undigested [bp] / digested [bp] to estimate their relative quantities.

### Quantitative PCR (qPCR)

DNA was extracted by homogenizing whole flies in 200µl Chelex solution at 5% concentration in distilled water. A total of 10µl of proteinase K (10mg/ml) was added per sample and the vials were incubated for 6 hours at 56°C. After digestion, vials were centrifuged at 16,200 x g for 3 minutes, and the supernatant was transferred to a new vial. DNA was stored at 4°C and freeze-thaw cycles were avoided when possible. Samples were run in duplicate using SYBR Green reagent on a StepOnePlus Real Time PCR machine (Applied Biosystems, USA). Technical repeats with >1 Ct difference were rerun or discarded. Each reaction contained 10µl Maxima SYBR Green/ROX qPCR Master Mix (2x), 1µl of forward primer, 1µl of reverse primer, 6µl nuclease-free water and 2µl of the DNA sample. Discriminant mitochondrial primers cycling conditions were 95°C for 20 seconds, followed by 40 cycles of: 95°C for 3 seconds and 55°C for 30 seconds (**Table S1**). Discriminant primers used linked base technology to increase specificity (Exiqon, Denmark). Melt curves and the fly control gene *rps17* were run on each plate to control for successful single-peak amplification and to control for DNA quality. All primers had >90% efficiency. For absolute quantification, each 96-well qPCR plate was analyzed using StepOne Software v2.2.2 (Applied Biosystems, USA) and Ct values were obtained by comparing each primer sample to a single standard curve of known concentration and using identical threshold and baseline levels for each primer target across plates. Standard curves were created by amplifying positive control samples using PCR, calculating DNA concentrations using QBIT, and then serially diluting the sample 1:10 with distilled water to create a 5-sample curve comprising known concentrations decreasing from 10pmol/ ml. Samples with Ct values over 30 were classed as negative, confirmed by our negative controls. As a result, the detection limit was 50 copies per sample for *rps17*, 115 for *CO3* (coxSNP_1) and 735 for *CO3* (coxSNP_2). DNA concentration was calculated for each sample by comparing the sample Ct values back to the equation of the standard curve. DNA copy numbers were then calculated by using the DNA product length to determine DNA concentration. To control for fly size and extraction efficiency, copy numbers for each sample were normalized using the fly housekeeping gene as a control, giving a final value in terms of mitochondrial genome copies to fly nuclear genome copy ratio. Since *rps17* is located on Muller element D, which in *D. paulistorum* is fused to Muller element A and part of the X chromosome, the copy numbers differ between males and females. Hence, male *rps17* numbers were doubled. We used ANOVA in R version 3.3.2 (R Core Team, 2016) when testing for significant changes in copy numbers of the α-NUMT and the real β mitochondrial genome between ages and sexes of different tissues. The data was log-transformed to increase normality.

### Generation of F1 hybrids

For mass-mating crosses in pools of virgin females and males from different *D. paulistorum* lines were obtained by collecting newly emerged flies within 2-3 h intervals to avoid any possible sib mating. Collected flies were kept separated for 5-7 days to ensure their virgin status. Crosses were set up in pools using 10-20 females and 20-30 males per vial. Flies were mated for one week and then transferred to new vials. F_1_ flies emerging from the vials were collected for DNA extraction. F_1_ individuals were used for RFLP-PCR assays.

### DNA extraction, genome sequencing and assembly

#### DNA extractions and Nanopore sequencing

Total DNA for Oxford Nanopore (ONT) sequencing of O11 was extracted from many adult flies of both sexes using the MagAttract HMW DNA Kit (Qiagen, Germany) with only minor modifications of the manufacturer’s instructions. Several similar independent DNA extractions were then pooled, cleaned and concentrated with the Ampure XP beads clean-up protocol (Beckman Coulter, USA). The Circulomics SRE XS kit (Pacific Biosciences, USA) was then used to eliminate short DNA fragments before sequencing. A DNA library was prepared using the Ligation Sequencing kit (Oxford Nanopore Technologies, UK) and sequenced on an R9.4 PromethION flowcell (FLO-PRO002) at the Uppsala Genome Center, Uppsala, Sweden. Basecalling was performed with Guppy 4.3.4 and the HAC model. A total of 1.4M reads were generated, with 19.54 Gb passed bases (**Table S3**).

#### D. paulistorum O11 genome assembly

Reads were subsampled with Filtlong v.0.2.1 (https://github.com/rrwick/Filtlong, last visited 18th March 2022) to lower the overall coverage of the reads to facilitate the assembly process. After testing different settings (**Table S3**), the O11 Nanopore reads were filtered to a coverage of 50X, giving priority to the quality of the reads (--target_bases 12500000000 –min_length 1000 – mean_q_weight 10). The whole genome assembly was built from the dataset of subsampled reads using NextDenovo v.2.5.0 (Hu et al., 2024; https://github.com/Nextomics/NextDenovo, last visited 18th March 2022) with a read cutoff of 1 Kb, and setting the genome size to 250 Mb. The genome assembly was then polished with both the same Nanopore reads and Illumina reads from Baião et al., 2023 (available via NCBI Bioproject PRJNA643793). First, Nanopore reads were mapped to the genome assembly with Minimap2 (Li, 2018) using the map-ont mode and bam files were sorted and indexed with samtools (Danecek et al., 2021). Then, the pipeline PEPPER-Margin-DeepVariant r0.4 (Shafin et al., 2021) was used to call variants. The variants were filtered with bcftools v1.15.1-21 (Danecek et al., 2021) using the following thresholds: QUAL>=30 && FMT/DP>=10 && FMT/GQ>=30 && FMT/VAF>=0.8, and remaining SNPs were corrected in the genome with bcftools v1.15.1-21 (Danecek et al., 2021). Illumina reads were aligned to the genome assembly using BWA mem v0.7.17-r1198-dirty (Li, 2013) and sorted with samtools v1.14 (Danecek et al., 2021), and two runs of Pilon v1.24 (Walker et al., 2014) were used for correction. BUSCO v5.2.2 with diptera_odb10 and augustus species fly was run to check genome assembly completeness, and Quast v5.0.2 (Mikheenko et al., 2018) was run with default settings to quantitatively assess the genome assembly stats. Scripts can be found in https://github.com/mmontonerin/Dpaulistorum_Mega-NUMT.

### Detection and annotation of α-NUMT copies in the genome assembly

The O11 β mt-genome (Baião et al., 2023) was used as a query when running BLASTn v2.12.0+ (Camacho et al., 2009) against the genome assembly. Hits over 500 bp long and with 90% identity to the mitochondrial sequence were inspected, and the 12 mt-genome copies in the Mega-NUMT region were identified on contig ctg000940. The position of each gene in the α-NUMT copy in the Mega-NUMT was obtained by using gene sequences from the *D. willistoni* mt-genome (GD-H4-1 line) as queries in a BLASTn search. Each inferred gene was subsequently checked manually in Artemis v.16 (Rutherford et al., 2000) to identify pseudogenes and adjust the position if needed. Scripts can be found in https://github.com/mmontonerin/Dpaulistorum_Mega-NUMT.

### Phylogenomic analysis of the α-NUMT copies

The 13 mitochondrial protein-coding genes from all α-NUMT copies and the genes from the real mt-genomes from several species of the willistoni group (see **Fig. 3**), previously published in Baião et al., 2023 (available via NCBI Bioproject PRJNA643793), were aligned using MAFFT v.7.407 (Katoh & Standley, 2013) and subsequently trimmed with trimAl v1.4.1 (Capella-Gutiérrez et al., 2009) using the setting -gt 0.1. The python script gene_stitcher.py (https://github.com/ballesterus/Utensils/blob/master/geneStitcher.py, last visited 12 March 2026) was used to concatenate the trimmed alignments. The concatenated alignment was used to infer a maximum likelihood phylogeny in IQTree2 v2.2.0 (Minh et al., 2020) with one partition per gene and the parameter settings -m MFP -b 100 -T AUTO. Scripts can be found in https://github.com/mmontonerin/Dpaulistorum_Mega-NUMT.

### Analyzing insertion sequences in α-NUMTs

Finally, we used a snakemake pipeline for repeat discovery and annotation in the genome assembly, which includes prediction of repeat content and transposable elements with RepeatModeler v1.0.8 (https://github.com/Dfam-consortium/RepeatModeler, last visited 12 March 2026). The pipeline also creates a database based on UniProt/Swissprot (The UniProt Consortium, 2025) protein dataset without transposable elements that initial predictions can be blasted against and removed if matching. By this, protein-coding genes in multiple copies are avoided in our final repeat library. The final repeat library was then used to mask the genome assembly with RepeatMasker v4.0.7 (https://github.com/Dfam-consortium/RepeatModeler, last visited 12 March 2026). Scripts can be found in https://github.com/mmontonerin/Dpaulistorum_Mega-NUMT.

### Collection of staged fly embryos for DNA extraction and DNA FISH

*D. paulistorum* embryos at early (0-30 min AEL), mid (3-4 hrs. AEL) and late (18-20 hrs. AEL) stages of development were collected on homemade medium (agar, molasses, cornmeal and yeast medium) in plastic Petri dishes (3 x 3 cm) using collection chambers at 25°C. Prior to collecting embryos, 1–2-week-old flies were kept in these chambers for 2-3 days to adapt to the new environment. Embryos were then collected by gently washing them off the plate onto a mesh with water. A subset of embryos was further processed for DNA FISH (see below). The rest was manually picked with a dissection needle and transferred immediately to an ice-cold lysis buffer (Qiagen, Germany) and processed for DNA extraction or stored at −20°C for no more than one week. In total, we collected 20 embryos for each sample with at least five biological replicates.

### Fluorescent in situ hybridization (FISH) for Mega-NUMT visualization

We performed DNA FISH on various fly tissues using both satellite and Mega-NUMT probes. The Mega-NUMT probe was generated by direct labeling via PCR with modified dNTPs (Alexa Fluor 488 dUTP; Jena Bioscience, Germany). We used a mixture of five mitochondrial loci (*CO1*, *CO2*, *ND1*, *ND2* and *ND4*) of *D. paulistorum*, cloned beforehand in pTZ57R/5 vector (Thermo Scientific, USA). For targeting the satellite sequence adjacent to the Mega-NUMT, we used a primary oligo probe coupled with a sec6 adaptor (satO11-CACACGCTCTCCGTCTTGGCCGTGGTCGATCAttttttttttTTTATAAAAATTAATACTGAGGACAAACTGAGGACT) and the secondary probe Sec6 coupled with Cy3 (Sec6-/5Cy3/aTGATCGACCACGGCCAAGACGGAGAGCGTGTGaa). The probe for the nuclear single copy gene *FOXO* gene was generated in the same way as the Mega-NUMT probe, but by only using three probes covering the *D. paulistorum FOXO* gene and labelled with Alexa Fluor 594 dUTP (Jena Bioscience, Germany).

To visualize Mega-NUMT in mitotically active neuroblasts of third instar larvae, we dissected their brains in PBS and incubated for 15 min in 0.5% sodium citrate. Then we fixed the brains for 15 min in 4% formaldehyde and 45% acetic acid before squashing them between the slide and coverslip. Squashed samples were immediately immersed in liquid nitrogen and transferred afterwards in 100% ethanol for 5 min. Then we air dried slides for at least 1 hour before proceeding to the hybridization. For the hybridization, we used 20 pmol of satellite probes coupled with 80 pmol of the sec6 probe and approximately 150 ng of Mega-NUMT probe in 50 μL of hybridization buffer (50% formamide, 10% dextran sulfate, 2xSSC). We heated slides for 5 min at 95°C to denature and incubated them overnight at 37°C in a humid chamber. We then washed the slides 3 times for 5 min with 4xSSCT and 3 times for 5 min with 0.1xSSC before mounting in SlowFade with DAPI and sealing with nail polish.

To visualize Mega-NUMT in mitotically active early embryos (0-3 h), staged embryos were collected as stated above, dechorionated in 50% bleach and fixed for 20 min in 3.7% paraformaldehyde in PBX (PBS and 0.15% Triton X-100) mixed with heptane (1:1 v/v). Then the embryos were postfixed in ice-cold methanol and further processed for FISH following a protocol by Gemkow et al., 1996 with slight modifications. Additionally, salivary glands and polytene chromosomes were used for Mega-NUMT visualization. The glands were dissected in PBS and fixed for 20 min in 3.7% paraformaldehyde in PBX (PBS and 0.15% Triton X-100). After washing three times in PBX, the tissues were kept in 70% ethanol until being used for DNA FISH. Polytene chromosomes were prepared according to (Pimpinelli et al., 2010) and processed for DNA FISH as described above.

Images were taken with an Olympus FW3000 laser scanning confocal microscope (Olympus Corporation, Japan) using a 60x objective and further processed with Photoshop CS6 (Adobe CS, USA), adjusting levels for each channel.

## Notes

### Competing Interest Statement

The authors have declared no competing interest.

### Summary of Updates

based on reviwers we slightly modified the earlier version

