## Supplement Material for "Evolutionary persistence of a highly prevalent multicopy mitochondrial-derived nuclear insertion (Mega-NUMT) in Neotropical *Drosophila* flies"

### Supplementary material

**Table S1. List of primer sets used for qPCR and RFLP-PCR.** Table lists the primer sequence and the according annealing temperature for each primer set used in PCR as well as the corresponding amplicon size.

| Locus | Primer name | 5'-3' primer sequence | Target | Tm | Fragment length | experiment |
|---|---|---|---|---|---|---|
| RPS17 | qRp 17 melwin 1 F | GCCGTCTGCGTCACTCC | Fly control gene | 60°C | 106bp | qPCR |
| RPS17 | qRp 17 melwil1_R | GCTTCAACATCTCCTTGGTATCG | Fly control gene | 60°C | 106bp | qPCR |
| CO3 | coxSNP_1 | GACATTTTGTTGATGTAGTT | alpha | 52°C | 121bp | discPCR |
| CO3 | coxSNP_2 | GACATTTTGTTGATGTAGTG | beta | 52°C | 121bp | discPCR |
| CO3 | coxSNP_uni | TAGACCTTATGATTGGAAGT | Universal reverse mitotype | 52°C | 121bp | discPCR |
| CO1 | CO1-F | TTTAATTTTACCTGGATTTGG | | 56°C | 788bp | RFLP-PCR |
| CO1 | CO1_wob-F | TTTAATTTTACCWGGATTTGG | | 56°C | 788bp | RFLP-PCR |
| CO1 | CO1_univ-R | ATTCAGAATATCTATGTTCAG | | 56°C | 788bp | RFLP-PCR |
| CO1 | CO1_wm-F | TTATTTACAGGATTAACTTTAAATGC | | 56°C | 705bp | RFLP-PCR |
| CO1 | CO1_wm-R | ACAGAAGGTTCATTAATTTCATC | | 56°C | 705bp | RFLP-PCR |
| CO2 | CO2-F | ATGTCTACATGAGCTAATTTAGGT | | 56°C | 323bp | RFLP-PCR |
| CO2 | CO2-R | TAACTTCAGTATCATTGGTGACC | | 56°C | 323bp | RFLP-PCR |
| CO3 | CO3-F | TATAATTCAATCTTATGTATTTGCAG | | 55°C | 841bp | RFLP-PCR |
| CO3 | CO3-R | GATTATTCCTTTATATTACAATTTACT | | 55°C | 841bp | RFLP-PCR |
| ND2 | ND2-F | CTTCYATTAATCATTTAGGATGAATATTAAGYGC | | 65°C | 357bp | RFLP-PCR |
| ND2 | ND2-R | AAGCTGAATAACAAATTCGTAARTAAAAAAATAAAG | | 65°C | 357bp | RFLP-PCR |

**Table S2. F-test statistics to test differences in relative copy number of β-mitochondria and α-NUMT across different parameters:** tissue, sex, age and combinations of those. The main results are shown in Figure 2. Df: degrees of freedom; sq: square.

| **Cox1-α** | **Df** | **Sum Sq** | **Mean Sq** | **F Value** | **P-value (>F)** |
|---|---|---|---|---|---|
| **Tissue** | 4 | 2.43 | 0.6070 | 1.728 | 0.150 |
| **Sex** | 1 | 0.77 | 0.7713 | 2.195 | 0.142 |
| **Age** | 1 | 0.00 | 0.0001 | 0.000 | 0.986 |
| **Tissue:Sex** | 4 | 0.10 | 0.0249 | 0.071 | 0.991 |
| **Tissue:Age** | 4 | 0.14 | 0.0343 | 0.097 | 0.983 |
| **Sex:Age** | 1 | 0.17 | 0.1660 | 0.472 | 0.493 |
| **Tissue:Sex:Age** | 4 | 1.05 | 0.2620 | 0.746 | 0.563 |
| **Residuals** | 100 | 35.13 | 0.3513 | | |
| **Cox2-β** | **Df** | **Sum Sq** | **Mean Sq** | **F Value** | **P-value (>F)** |
| **Tissue** | 4 | 26.525 | 6.631 | 27.694 | 1.76e-15*** |
| **Sex** | 1 | 1.129 | 1.129 | 4.714 | 0.0323* |
| **Age** | 1 | 0.011 | 0.011 | 0.044 | 0.8343 |
| **Tissue:Sex** | 4 | 1.145 | 0.286 | 1.195 | 0.3175 |
| **Tissue:Age** | 4 | 0.447 | 0.112 | 0.467 | 0.7598 |
| **Sex:Age** | 1 | 1.596 | 1.596 | 6.666 | 0.0113* |
| **Tissue:Sex:Age** | 4 | 1.269 | 0.317 | 1.325 | 0.2660 |
| **Residuals** | 100 | 23.945 | 0.239 | | |

**Table S3.** Genome assembly information across different assembly methods. Unfiltered refers to raw reads being used directly in Nextdenovo to produce the genome assembly. Phred filters refer to the priority of quality of the reads when subsampling reads to the marked coverage (X40, X50). Length sets priority in read length when subsampling reads. Two variations (noted with "+ assembly ref") included the unfiltered assembly as reference for the subsampling. The second column shows how many reads were left after subsampling, and the third column indicates the total number of base pairs in the dataset assembled. Quast was used to assess assembly size, contigs, N50 and the size of the largest contig. BUSCO was used to assess genome completeness based on single-copy genes.

| O11 | Reads | bp | Coverage | Assembly size | Contigs | N50 | BUSCO | Largest contig |
|---|---|---|---|---|---|---|---|---|
| unfiltered | 1 376 601 | 19 454 247 303 | 77 | 251 478 094 | 188 | 28 092 762 | C:97.3%[S:96.3%,D:1.0%],F:1.1%,M:1.6%,n:3285 | 35 123 609 |
| Phred X40 | 641 549 | 10 000 005 368 | 40 | 224 185 510 | 96 | 16 811 753 | C:97.2%[S:96.4%,D:0.8%],F:1.1%,M:1.7%,n:3285 | 29 505 846 |
| Phred X40 + assembly ref | 472 844 | 10 000 013 032 | 40 | 225 554 080 | 107 | 15 921 554 | C:97.5%[S:96.6%,D:0.9%],F:1.0%,M:1.5%,n:3285 | 31 708 369 |
| Phred X50 | 827 899 | 12 500 020 744 | 50 | 231 244 016 | 111 | 18 692 465 | C:97.6%[S:96.7%,D:0.9%],F:1.2%,M:1.2%,n:3285 | 33 659 176 |
| Length X40 | 312 151 | 10 000 014 293 | 40 | 235 601 215 | 126 | 13 106 850 | C:97.1%[S:96.3%,D:0.8%],F:1.1%,M:1.8%,n:3285 | 29 628 043 |
| Length X40 + assembly ref | 296 871 | 10 000 014 326 | 40 | 231 875 968 | 113 | 25 388 691 | C:97.2%[S:96.3%,D:0.9%],F:1.1%,M:1.7%,n:3285 | 37 599 843 |
| Length X50 | 827 899 | 12 500 012 596 | 50 | 239 343 741 | 128 | 16 562 743 | C:97.2%[S:96.4%,D:0.8%],F:0.9%,M:1.9%,n:3285 | 39 187 662 |

**Table S4. Polishing iterations of the genome assembly**, accounting for the number of fixes that each method has produced, and the final BUSCO score. They are ordered in the order in which they were done over the assembly Pepper -> Pilon1 -> Pilon2 -> Pilon3. The assembly polished by two rounds of pilon was selected as the optimal genome assembly, to avoid introducing errors.

| Polishing method | Insertions | Deletions | SNPs | Total fix | BUSCO |
|---|---|---|---|---|---|
| PEPPER | 68 | 1719 | 4071 | 4071 | C:97.6%[S:96.7%,D:0.9%],F:1.2%,M:1.2%,n:3285 |
| Pilon 1 | 139108 | 37585 | 0 | 176693 | C:99.0%[S:98.1%,D:0.9%],F:0.3%,M:0.7%,n:3285 |
| Pilon 2 | 1261 | 1719 | 0 | 2980 | C:99.0%[S:98.1%,D:0.9%],F:0.3%,M:0.7%,n:3285 |
| Pilon 3 | 176 | 535 | 0 | 711 | C:99.0%[S:98.1%,D:0.9%],F:0.3%,M:0.7%,n:3285 |

**Table S5. Genes and pseudogenes of α-NUMTs in the genome assembly of *D. paulistorum* O11.** Each row represents one NUMT copy found in the genome assembly, while the columns represent the 13 mitochondrial genes of the *D. paulistorum* mitochondrial genome (listed below). Green = complete gene, yellow = pseudogene, grey = absence of the gene. *ND2*: NADH dehydrogenase subunit2, *CO1*: cytochrome c oxidase subunit I, *CO2*: cytochrome c oxidase subunit II, *ATP8*: ATP synthase F0 subunit 8, *ATP6*: ATP synthase F0 subunit 6, *CO3*: cytochrome c oxidase subunit III, *ND3*: NADH dehydrogenase subunit 3, *ND5*: NADH dehydrogenase subunit 5, *ND4*: NADH dehydrogenase subunit 4, *ND4L*: NADH dehydrogenase subunit 4L, *ND6*: ND6 dehydrogenase subunit 6, *cytB*: Cytochrome b, *ND1*: NADH dehydrogenase subunit 1.

| | *ND2* | *CO1* | *CO2* | *ATP8* | *ATP6* | *CO3* | *ND3* | *ND5* | *ND4* | *ND4L* | *ND6* | *cytB* | *ND1* |
|---|---|---|---|---|---|---|---|---|---|---|---|---|---|
| NUMT 1 | | | | | | | | | | | | | |
| NUMT 2 | | | | | | | | | | | | | |
| NUMT 3 | | | | | | | | | | | | | |
| NUMT 4 | | | | | | | | | | | | | |
| NUMT 5 | | | | | | | | | | | | | |
| NUMT 6 | | | | | | | | | | | | | |
| NUMT 7 | | | | | | | | | | | | | |
| NUMT 8 | | | | | | | | | | | | | |
| NUMT 9 | | | | | | | | | | | | | |
| NUMT 10 | | | | | | | | | | | | | |
| NUMT 11 | | | | | | | | | | | | | |
| NUMT 12 | | | | | | | | | | | | | |

**Table S6. List of fly lines used in the study belonging to willistoni group species.** Abbreviation: NDSSC - National Drosophila Species Stock Center

| Species | Subspecies | Line | Locality | Source/Collector | Date |
|---|---|---|---|---|---|
| *D. equinoxialis* | | FS | Fort Sherman, Colon, Panama | Egon Bartel | 2002 |
| *D. insularis* | | MASS-B-SL | St Lucia | Jeff Powell | 2006 |
| *D. tropicalis* | | SS | San Salvador, El Salvador | NDSSC: 14030-0801.01/ William Heed | 1955 |
| *D. willistoni* | | APA8-2 | Veracruz, México | Joana Silva | 1998 |
| | | FG168 | Saül, French Guiana, France | Wolfgang Miller & Aurélie Hua-Van | 2014 |
| *D. paulistorum* | *Amazonian* | A28 | Belém, PA, Brazil | Lee Ehrman | 1952 |
| | | FG572 | Saül, French Guiana, France | Wolfgang Miller & Aurélie Hua-Van | 2015 |
| | | FG16 | Saül, French Guiana, France | Wolfgang Miller & Aurélie Hua-Van | 2018 |
| | | FG18 | Saül, French Guiana, France | Wolfgang Miller & Aurélie Hua-Van | 2018 |
| | *Andean-Brazilian* | MS | Mesitas, Colombia | Lee Ehrman | 1962 |
| | | Yellow | Copan, Honduras | NDSSC: 14030-0771.02 | 1960 |
| | *Centro american* | C2 | Lancetilla, Honduras | Lee Ehrman | 1954 |
| | | White | Lancetilla, Honduras | NDSSC: 14030-0771.03 | 1960 |
| | *Interior* | L1 | Llanos, Colombia | Lee Ehrman | 1958 |
| | *Orinocan* | O11 | Georgetown, Guyana | Lee Ehrman | 1957 |
| | | RP | Ribeirão Preto, SP, Brazil | Ana Laurer Garcia | 1995 |
| | | POA1 | Porto Alegre, RS, Brazil | Ana Laurer Garcia | 2003 |
| | | TP37 | Pousada Triunfo, PE, Brazil | Claudia Rohde & Ana Laurer Garcia | 2009 |
| | | FG103 | Saül, French Guiana, France | Wolfgang Miller & Aurélie Hua-Van | 2014 |
| | | FG111 | Saül, French Guiana, France | Wolfgang Miller & Aurélie Hua-Van | 2014 |
| | | FG295 | Saül, French Guiana, France | Wolfgang Miller & Aurélie Hua-Van | 2014 |
| | *Transitional* | SM | Santa Marta, Colombia | Lee Ehrman | 1956 |
| *D. nebulosa* | | NEB | Palmira, Colombia | NDSSC: 14030-0761.00/ William Heed | 1955 |

**Table S7. Mega-NUMT prevalence in *D. paulistorum* from recent collections in French Guiana**

| Line Origin[1] | Collection Date | Latitude | Longitude | Mega-NUMT[2] | Prevalence |
|---|---|---|---|---|---|
| **Long-term stocks from 1960s** | *see* Table 1 and Table S6 | | | **4/9** | **0.44** |
| **French Guiana, F** | | | | | |
| CNRS Camp Inselberg | June 2015 | 04˚05'16.73" N | 52˚40'47.31"W | 8/23 | 0.35 |
| CNRS Camp Inselberg | Aug. 2019 | 04˚05'16.73" N | 52˚40'47.31"W | 4/8 | 0.50 |
| CNRS Camp Pararé | May 2014 | 04˚02'16.28" N | 52˚40'22.55"W | 2/5 | 0.40 |
| Route to Kaw | Sept. 2024 | 04˚33'34.27" N | 52˚12'26.11"W | 6/14 | 0.43 |
| Saül, Chez Lulu | May 2014 | 03˚37'16.35" N | 53˚12'41.62"W | 8/15 | 0.53 |
| Saül, Chez Lulu | August 2019 | 03˚37'16.35" N | 53˚12'41.62"W | 4/11 | 0.36 |
| Maripasoula, Le terminus | May 2014 | 03˚38'09.18" N | 54˚01'44.58"W | 2/4 | 0.50 |
| **French Guiana total:** | 2014-2024 | | | **34/80** | **0.43** |

[1] Geographic origin of the established isofemale line.
[2] Number of isolines carrying a Mega-NUMT and number of total isolines tested are based on diagnostic double-peaks in chromatograms of direct *CO1* Sanger sequencing and cloning. For field samples from French Guiana, Mega-NUMT-typing was performed on multi-fly DNA from established isofemale lines at generation F1 and F2.

**Table S8. Mega-NUMT-typing of the 80 isofemale lines collected between 2014 and 2024 at different localities in French Guiana (summarized in Figure 7)**. Mega-NUMT-typing was performed by *CO1* direct Sanger sequencing and/or quantitative RFLP-PCR for abundance and prevalence for samples with available data sets. After more than 10 years under lab conditions (highlighted in gray and tested again in 2026), only 7 out of the 16 originally MN-positively typed at F1/F2 lines have maintained the Mega-NUMT (43.75%), dropping from initial 51.6% (16/31) to 22.6% (7/31) under lab conditions.

| French Guiana Iso-Line | Sampling Year | *D. paulistorum* Spp | Mito-type | α-NUMT detected in F1 by Sanger | α-NUMT detected in 2026 by Sanger | α-NUMT Abundance (%)[#] f/m | α-NUMT Prevalence |
|---|---|---|---|---|---|---|---|
| **Maripousala** | | | | | | | |
| FG127 | 2014 | AM | ß1 | - | - | 0 | |
| FG128 | 2014 | AM | ß1 | - | - | 0 | |
| FG219 | 2014 | AM | ß1 | Yes | - | ND | ND |
| FG238 | 2014 | AM | ß1 | Yes | NA | ND | ND |
| **CNRS Camp Pararé** | | | | | | | |
| FG248 | 2014 | AM | ß1 | - | - | 0 | |
| FG250 | 2014 | AM | ß1 | Yes | NA | 57/62.7 | ND |
| FG254 | 2014 | AM | ß1 | **Yes** | **Yes** | 27.6/66.0 | ND |
| FG256 | 2014 | AM | ß1 | - | - | 0 | |
| FG259 | 2014 | AM | ß1 | - | - | 0 | |
| **Saül** | | | | | | | |
| FG82 | 2014 | OR | ß2 | - | NA | 0 | 0% (10/10) |
| FG89 | 2014 | AM | ß1 | Yes | - | ND | 0% (0/40) |
| FG97 | 2014 | OR | ß2 | - | NA | 0 | 0% (10/10) |
| FG103 | 2014 | OR | ß2 | Yes | - | ND | 0% (0/40) |
| FG104 | 2014 | OR | ß2 | - | - | 0 | 0% (10/10) |
| FG105 | 2014 | OR | ß2 | Yes | - | 18.6/24.2 | 0% (19/20) |
| FG109 | 2014 | OR | ß2 | Yes | NA | ND | ND |
| FG111 | 2014 | OR | ß2 | **Yes** | **Yes** | 10.3/11.8 | 55% (22/40) |
| FG113 | 2014 | OR | ß2 | **Yes** | **Yes** | 6.8/12 | 80% (16/20) |
| FG160 | 2014 | AM | ß1 | - | NA | 0 | 0% (10/10) |
| FG169 | 2014 | AM | ß1 | **Yes** | **Yes** | 58.6/52.6 | 15% (3/20) |
| FG186 | 2014 | AM | ß1 | Yes | - | 42.4/45.7 | 35% (7/20) |
| FG217 | 2014 | AM | ß1 | - | - | 0 | 0% (10/10) |
| FG271 | 2014 | AM | ß1 | - | - | 0 | 0% (10/10) |
| FG295 | 2014 | OR | ß2 | - | - | 0 | 0% (0/40) |
| **Saül** | | | | | | | |
| FG-MS1 | 2019 | AM | ß1 | - | ND | 0 | |
| FG-MS2 | 2019 | AM | ß1 | - | ND | 0 | |
| FG-WS1 | 2019 | AM | ß1 | - | ND | 0 | |
| FG-WS2 | 2019 | AM | ß1 | - | ND | 0 | |

| | | | | | | | |
|---|---|---|---|---|---|---|---|
| FG-WS3 | 2019 | AM | ß1 | Yes | ND | 50.8/49.7 | 100% (20/20) |
| FG-WS6 | 2019 | OR | ß2 | Yes | ND | NA | NA |
| FG-WS7 | 2019 | OR | ß2 | Yes | ND | 49.7/54.5 | 50% (10/20) |
| FG-WS12 | 2019 | OR | ß2 | Yes | ND | NA | NA |
| FG-WB2 | 2019 | AM | ß1 | - | ND | 0 | |
| FG-WB3 | 2019 | AM | ß1 | - | ND | 0 | |
| FG-WB4 | 2019 | AM | ß1 | - | ND | 0 | |
| **Camp CNRS Inselberg** | | | | | | | |
| FG508 | 2015 | AM | ß1 | Yes | NA | 31/25.2 | 10% (2/20) |
| FG509 | 2015 | AM | ß1 | Yes | - | 51.1/69.8 | 50% (10/20) |
| FG526 | 2015 | AM | ß1 | - | NA | 0 | |
| FG528 | 2015 | AM | ß1 | - | NA | 0 | |
| FG529 | 2015 | AM | ß1 | **Yes** | **Yes** | 47.2/54.2 | 35% (7/20) |
| FG546 | 2015 | AM | ß1 | Yes | - | NA | NA |
| FG547 | 2015 | AM | ß1 | - | NA | 0 | |
| FG548 | 2015 | AM | ß1 | - | NA | 0 | |
| FG549 | 2015 | AM | ß1 | - | NA | 0 | |
| FG550 | 2015 | AM | ß1 | - | - | 0 | |
| FG559 | 2015 | AM | ß1 | **Yes** | **Yes** | 39.7/52.4 | 80% (16/20) |
| FG569 | 2015 | AM | ß1 | Yes | NA | NA | NA |
| FG570 | 2015 | AM | ß1 | Yes | - | 46.1/44.6 | 50% (10/20) |
| FG572 | 2015 | AM | ß1 | **Yes** | **Yes** | 47.3/68.7 | 95% (19/20) |
| FG585 | 2015 | AM | ß1 | - | NA | 0 | |
| FG594 | 2015 | AM | ß1 | - | NA | 0 | |
| FG601 | 2015 | AM | ß1 | - | - | 0 | |
| FG602 | 2015 | AM | ß1 | - | - | 0 | |
| FG609 | 2015 | AM | ß1 | - | - | 0 | |
| FG614 | 2015 | AM | ß1 | - | NA | 0 | |
| FG617 | 2015 | AM | ß1 | - | - | 0 | |
| FG618 | 2015 | AM | ß1 | - | - | 0 | |
| FG620 | 2015 | AM | ß1 | - | - | 0 | |
| **Camp CNRS Inselberg** | | | | | | | |
| FG-WI10 | 2019 | AM | ß1 | - | ND | 0 | |
| FG-WI14 | 2019 | AM | ß1 | Yes | ND | 58.97/64.19 | 40% (8/20) |
| FG-WI17 | 2019 | AM | ß1 | Yes | ND | 61.39/64.49 | 80% (16/20) |
| FG-WI21 | 2019 | AM | ß1 | - | ND | 0 | |
| FG-WI24 | 2019 | AM | ß1 | - | ND | 0 | |
| FG-WIP1 | 2019 | AM | ß1 | Yes | ND | 41.04/71.59 | 75% (15/20) |
| FG-WIP2 | 2019 | AM | ß1 | - | ND | 0 | |
| FG-WIP3 | 2019 | AM | ß1 | Yes | ND | 34.72/55.55 | 80% (16/20) |
| **Road to Kaw** | | | | | | | |
| FG-A4 | 2024 | AM | ß1 | - | ND | 0 | |
| FG-A5 | 2024 | AM | ß1 | - | ND | 0 | |
| FG-A8 | 2024 | AM | ß1 | - | ND | 0 | |
| FG-A11 | 2024 | AM | ß1 | - | ND | 0 | |

| | | | | | | | |
|---|---|---|---|---|---|---|---|
| FG-A12 | 2024 | AM | ß1 | - | ND | 0 | |
| FG-A15.1 | 2024 | AM | ß1 | Yes | ND | NA | NA |
| FG-A16.1 | 2024 | AM | ß1 | Yes | ND | NA | NA |
| FG-A14 | 2024 | AM | ß1 | Yes | ND | NA | NA |
| FG-A21.1 | 2024 | AM | ß1 | Yes | ND | NA | NA |
| FG-A22 | 2024 | AM | ß1 | - | ND | 0 | |
| FG-A24 | 2024 | AM | ß1 | - | ND | 0 | |
| FG-A28 | 2024 | AM | ß1 | - | ND | 0 | |
| FG-A37 | 2024 | AM | ß1 | Yes | ND | NA | NA |
| FG-AB3 | 2024 | AM | ß1 | Yes | ND | NA | NA |

Abbreviations: NA: not available, because the line was lost over the last 10 years in the laboratory; ND: not determined; Yes: presence of the α-NUMT detected by the presence of double peaks in *CO1* chromatograms, and/or by the presence of the diagnostic *Hin*dIII RFLP restriction site. Gray backgrounded lines: Isolines that were maintained for more than 10 years under laboratory conditions and recently *CO1*-barcoded by direct Sanger sequencing in pools of 4 females and 6 males (n=31). Bold: Lines that have maintained in 2026 their Mega-NUMTs under laboratory conditions, like drift and artificial selection. (-) Absence of the MN in direct Sanger reads.

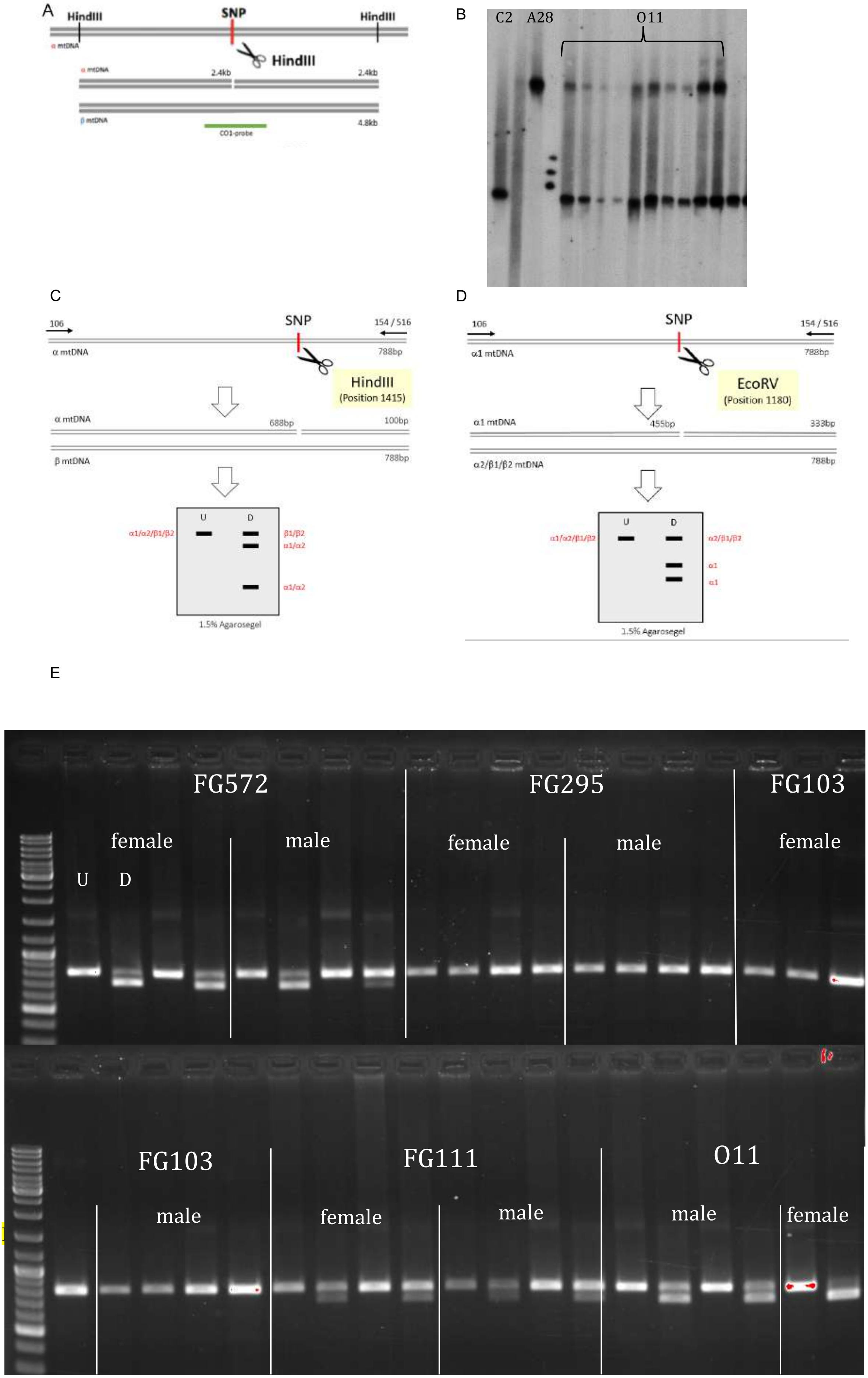
A
HindIII
SNP
HindIII
HindIII
2.4kb
2.4kb
4.8kb
B
C2
A28
O11
C
106
SNP
154 / 516
α mtDNA
788bp
HindIII
(Position 1415)
α mtDNA
688bp
100bp
β mtDNA
788bp
U
D
α1/α2/β1/β2
β1/β2
α1/α2
α1/α2
1.5% Agarosegel
D
106
SNP
154 / 516
α1 mtDNA
788bp
EcoRV
(Position 1180)
α1 mtDNA
455bp
333bp
α2/β1/β2 mtDNA
788bp
U
D
α1/α2/β1/β2
α2/β1/β2
α1
α1
1.5% Agarosegel
E
FG572
female
male
U
D
FG295
female
male
FG103
female
FG103
male
FG111
female
male
O11
male
female

**F**

Single Flies line FG572
(α2, ß1) – *COI, Hind*III

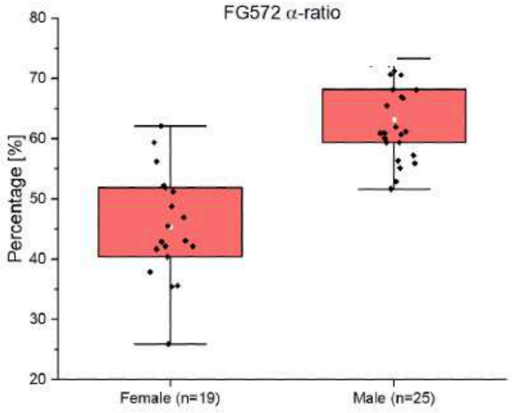

| COI | α - Mean | SD | Transmission |
|---|---|---|---|
| female (n=19) | 45.34 | 8.91 | 95% |
| male (n=25) | 63.19 | 6.54 | 83% |

Single Flies MS
(α1, α2) – *COI, Eco*RV

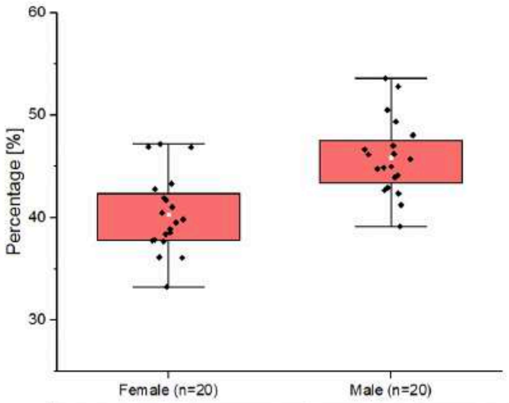

| COI | α2 - Mean | SD | Transmission |
|---|---|---|---|
| female (n=20) | 40.31 | 3.76 | 100% |
| male (n=20) | 45.86 | 3.67 | 100% |

Single Flies FG16
(α2, ß1) – *COI, Hind*III

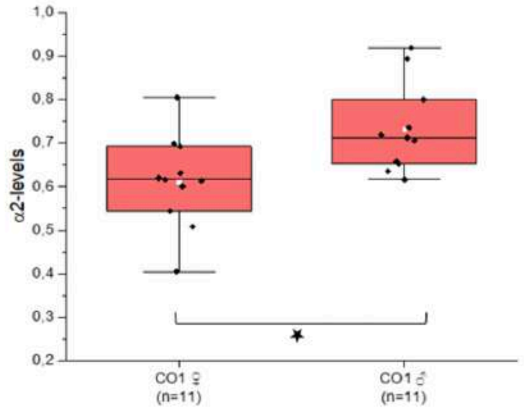

| COI | α2 - Mean | SD | Transmission |
|---|---|---|---|
| female (n=20) | 57.2 | 8.6 | 55% |
| male (n=20) | 66.5 | 57.2 | 55% |

Single Flies FG111
(α2, ß1) – *COI, Hind*III

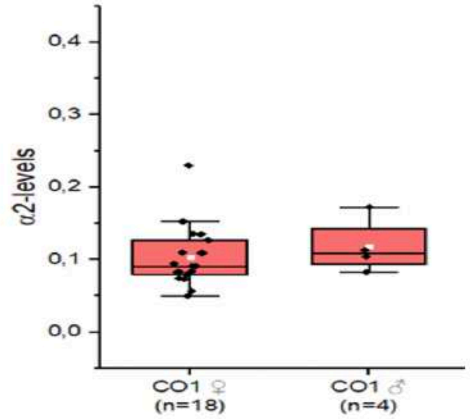

| COI | α2 - Mean | SD | Transmission |
|---|---|---|---|
| female (n=20) | 10.3 | 4.0 | 90% |
| male (n=20) | 11.8 | 3.2 | 20% |

**Figure S1. Schematic illustration of Restriction Fragment Length Polymorphism (RFLP) for differentiating α from β mitotypes in *D. paulistorum*. (A)** α-mitotype-specific RFLP-Southern blot. Whole fly DNA was fully digested with *Hin*dIII-enzyme, since only the α-mitotype displays the diagnostic SNP within the *CO1* restriction site (scissor), generating two 2.4 kb fragments, whereas the β-mitotype, lacking this restriction site at this position, thereby retains the 4.8 kb fragments. The internal *CO1*-probe of 788 bp cloned in a plasmid (green) covering the diagnostic SNP was used for their detection by Southern hybridization (**B**) Independent *CO1* Southern blot on *Hin*dIII digested DNA of C2 and A28 (each derived from a pool of 10 females) and 10 single fly O11 females probed with *CO1*. Equal DNA concentrations of app. 1μg was loaded per lane. Signal intensity differences between O11 samples represent quantitative differences in total DNA extracted from single flies. **(C)** Schematic description of the α-diagnostic RFLP-PCR assay. Similar to RFLP-Southern blot, the same diagnostic *Hin*dIII SNP (scissor) was used. PCR from total DNA with the consensus primer 106 and 154 (arrows) in *CO1* give rise to a 788 bp fragment in both mitotypes (U). After digestion (D), the β-mitotype remains undigested due to the absence of the *Hin*dIII site (upper 788 bp band in D), but the α-mitotype is represented by two bands of 688 bp and 100 bp length. **(D)** Schematic description of the α1-diagnostic RFLP-PCR assay. Here, diagnostic *Eco*RI SNP (scissor) was used. PCR from total DNA with the consensus primer 106 and 154 (arrows) in *CO1* give rise to a 788 bp fragment in both mitotypes (U). After digestion (D), the β-mitotype remains undigested due to the absence of the *Eco*RI site (upper 788 bp band in D), but the α1-mitotype is represented by two bands of 455 bp and 333 bp length. (**E**) Representative genotyping of the Mega-NUMT positive lines shown in **Table 1** by two randomly picked females and males per line, *CO1* RFLP/*Hin*dIII*,* undigested (U) and digested (D). (**F**) Mega-NUMT genotyping via RFLP-PCR of four independent representative *D. paulistorum* isolines. For details see Material and Methods.

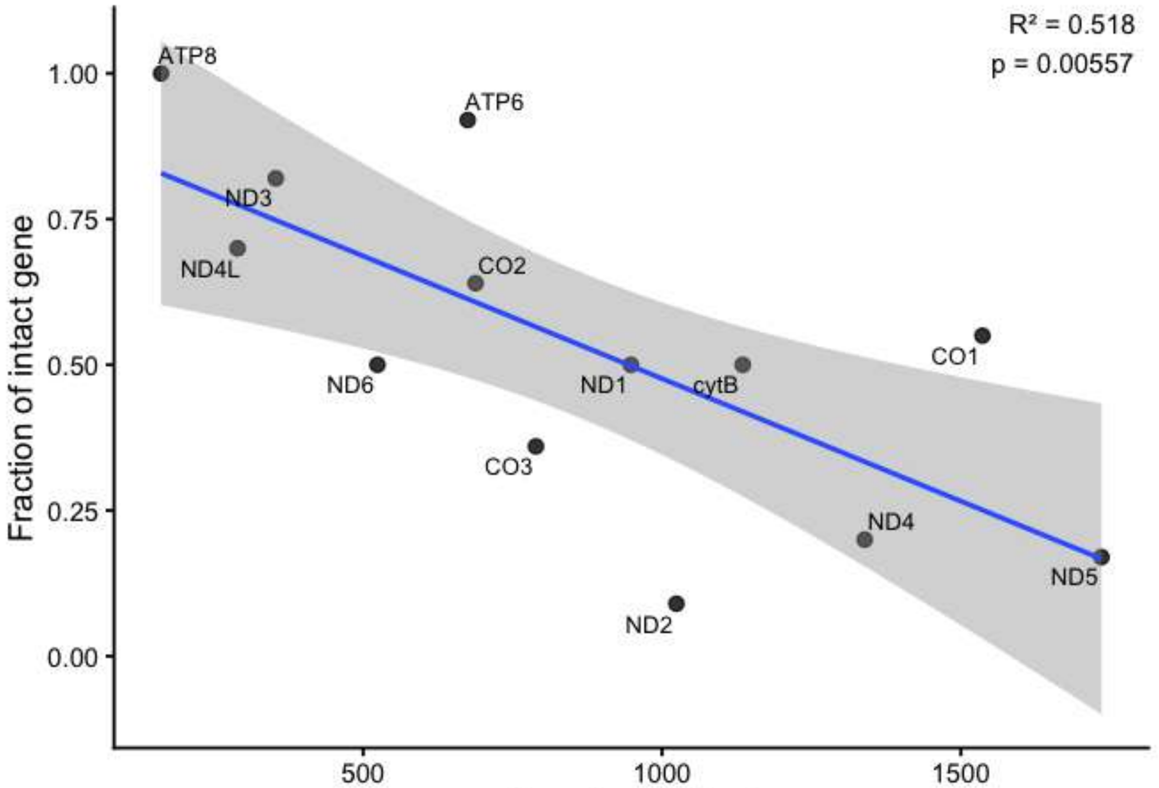

**Figure S2. Relationship between gene length and fraction of intact gene across the 12 alpha-NUMT copies.** Scatterplot showing gene length (bp) versus fraction of intact gene for individual genes. The fraction of intact genes was calculated as the number of copies with a pseudogenizing mutation divided by the number of copies present for each gene (see **Table S5**), and the gene length is the length of the intact gene. The solid line represents the linear regression fit, with the shaded area indicating the 95% confidence interval. $R^2$ and p-value are from a linear regression model testing the association between gene length and fraction intact.

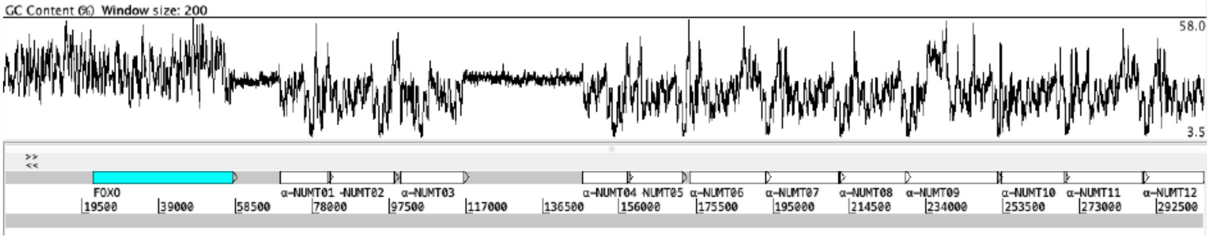

**Figure S3. GC content (%) across the end region of contig000940.** First base represents base 7.300.000 of the contig. Sliding window size for GC content values is 200. The nuclear gene *FOXO* is marked in blue. The α-NUMT copies are marked in white. GC values in the satellite sequences (between FOXO and α-NUMT01, and between α-NUMT03 and α-NUMT04) are low, with an average of 29.53% and 30.44% GC content, respectively, and very constant throughout the sequence. The average GC content throughout the nuclear genome assembly is 37.76 %, and the average GC content across the 12 α-NUMTs is 24.63%.

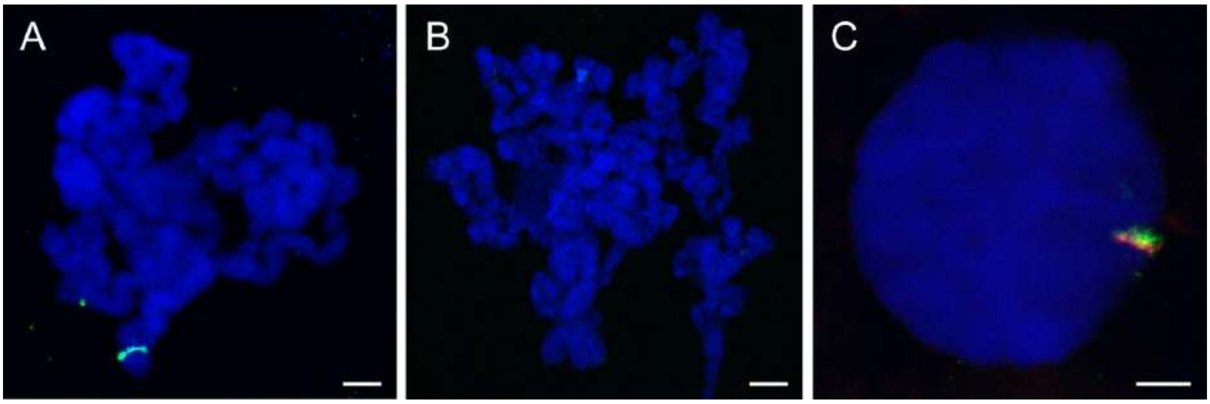

**Figure S4. Localization of the Mega-NUMT in *D. paulistorum* O11 line chromosomes 3 using DNA FISH with mitochondria-specific probes. (A)** Mega-NUMT band (green) on polytene chromosome 3 from larval salivary glands of the O11 line. Note the position of the band at the end of the chromosome arm. **(B)** The Mega-NUMT is absent in the FG295 line used as a control in the study. **(C)** Co-localization of the mega-NUMT (green) with the *FOXO* gene (red) in the salivary gland nucleus.

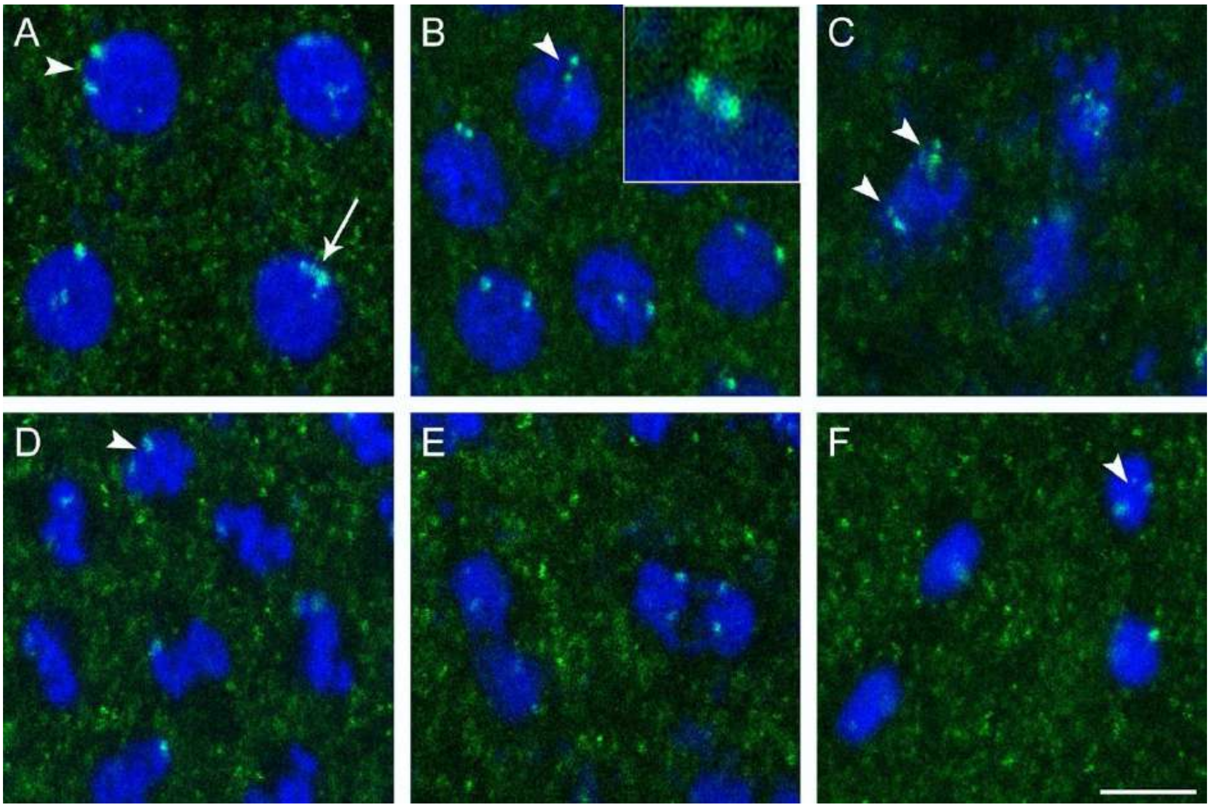

**Figure S5. Mega-NUMT dynamics in blastodermal embryo mitosis.** In mitotic nuclei at interphase **(A)**, prophase **(B)**, prometaphase **(C)**, metaphase **(D)** and telophase **(F),** the Mega-NUMT is observed mostly as a two-dot signal (green, arrowheads) with variegating distances between the dots. In some cases, the dots are fused into one big dot (green, arrow). In anaphase **(E)**, the Mega-NUMT signal is represented as 4 dots (2 dots per each segregating set of chromosomes). DNA is stained with DAPI (blue). Scale bar: 5 μm.

A

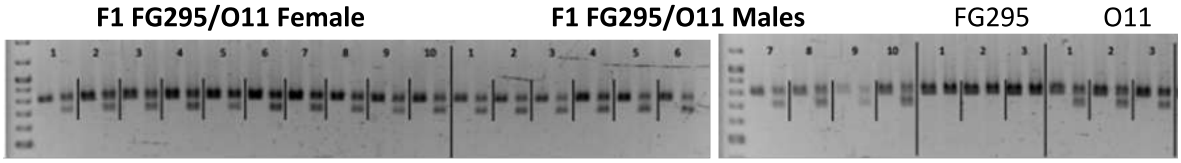

B

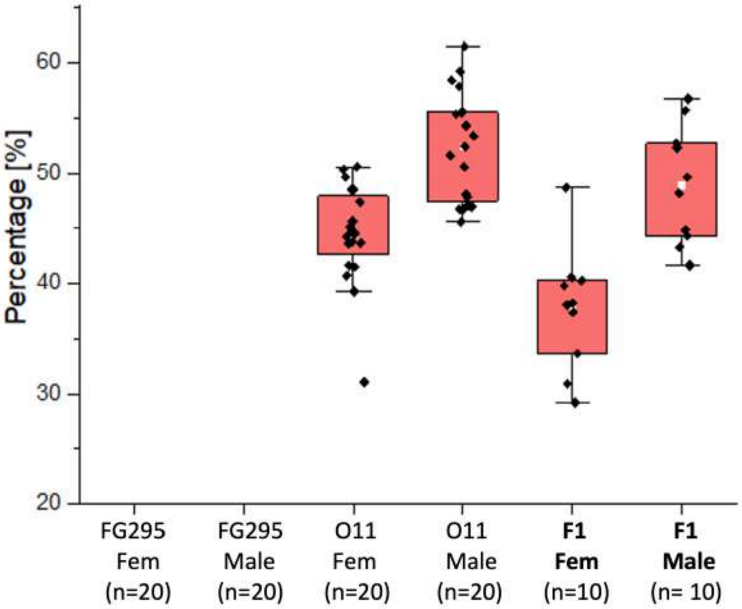

C

| *D. paulistorum* line | Mean MN abundance | SD | Transmission rate |
|---|---|---|---|
| O11 females (n=20) | 44.48 | 4.49 | 100% |
| O11 males (n=20) | 52.47 | 4.84 | 100% |
| FG295 females (n=20) | 0 | 0 | 0% |
| FG295 males (n=20) | 0 | 0 | 0% |
| **F1 females (n=10)** | **37.67** | **5.54** | **100%** |
| **F1 males (n=10)** | **48.94** | **5.35** | **100%** |

**Figure S6. Perfect transmission of the *Dpau* Mega-NUMT (MN) in F1 single fly hybrids (bold) of FG295 (MN-negative) and O11 (MN-positive) parents.** (**A**) *CO1* RFLP/*Hin*dIII-PCR of female and male F1 hybrids (bold) and their parental lines (for technical details see **Fig. S1**). (**B**) MN signal quantification of the *Hin*dIII sensitive band in individual flies. (**C**) Perfect transmission rate in F1s (n=20) confirming Mega-NUMT homozygosity in the O11 line of *D. paulistorum*.

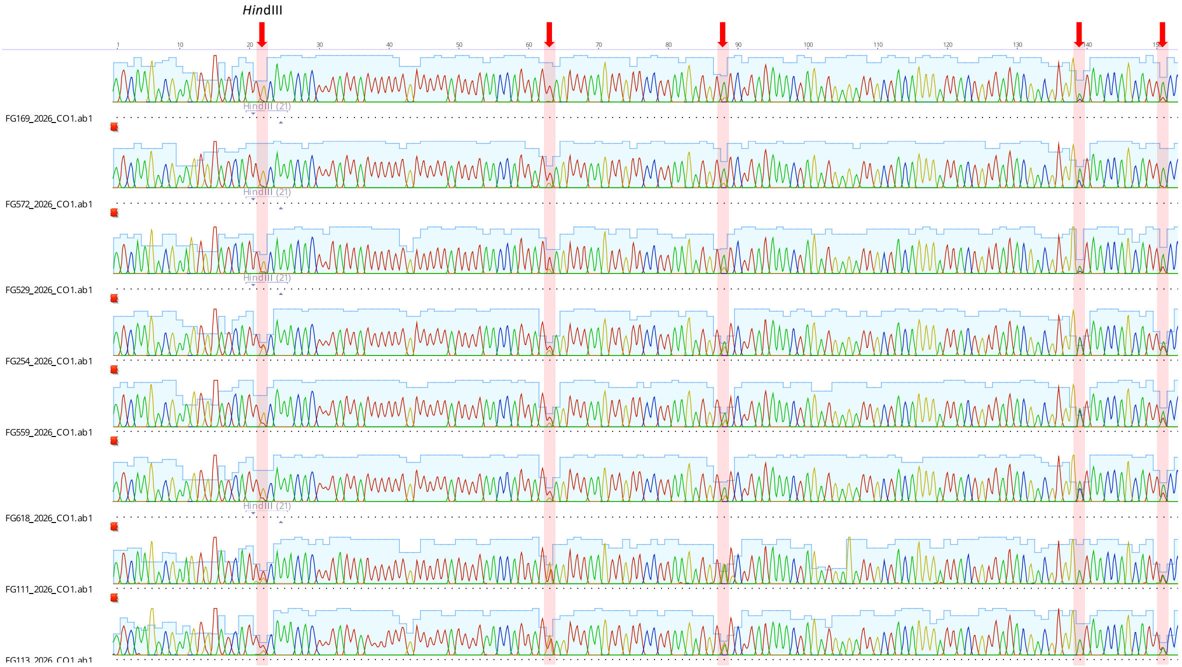

**Figure S7. Example of aligned *CO1* chromatograms of Mega-NUMT carrying candidate isolines from French Guiana by direct Sanger re-sequencing in 2026**. Polymorphic sites in *CO1* indicating the presence of the Mega-NUMT of *D. paulistorum spp.* by double-peeks (red arrow and highlighted in pink) and their diagnostic *Hin*dIII restriction site of isolines that have persisted under lab conditions from 2014 to 2026.
